# Defining ecologically realistic biodiversity offset multipliers with the Response-based Habitat Hectare Assessment of Biodiversity Gains (REHAB)

**DOI:** 10.64898/2026.01.26.701764

**Authors:** Joel Jalkanen, Eini Nieminen, Aapo Ahola, Emma Luoma, Minna Pekkonen, Panu Halme, Janne S. Kotiaho, Heini Kujala

## Abstract

In biodiversity offsetting, balancing biodiversity losses with gains can be achieved by using multipliers that define the ratio between the magnitude of biodiversity loss and the area required to deliver equivalent biodiversity gains. Although there is broad scientific consensus that multipliers should be calibrated to deliver no net loss or a net gain for biodiversity, they are often applied without quantitative assessment of the ecological outcomes of offset actions. Here we operationalise the ‘Response-based Habitat Hectare Assessment of Biodiversity Gains’ (REHAB), a framework where multipliers are informed by an understanding of habitat-specific ecological responses to offset action. To support Finland’s national biodiversity offsetting scheme, we harnessed the knowledge of 111 experts to compile ecological attributes and condition matrices for all 388 Finnish habitat types and derive 346 offset action multipliers that represent ecological response functions for 216 habitat type-specific offset actions including restoration, management and passive recovery. Our analysis reveals substantial variation in response-functions, resulting in offset multipliers between 1.3–4,000 across offset actions and habitat types. We find that the fixed multipliers commonly used in offset schemes would result in net loss in 60% of the cases if action– habitat specific responses were not considered. This variability underscores that fixed multipliers cannot deliver reliable biodiversity outcomes and should be avoided in offsetting schemes. The REHAB framework has already been integrated into Finland’s national offsetting policy. Other potential areas of application include informing ecosystem restoration planning and assessing biodiversity gains linked to credit issuance in emerging nature-credit markets.

## 1 Introduction

Biodiversity offsetting (hereafter offsetting) holds a promise that biodiversity losses can be compensated by achieving equivalent biodiversity gains elsewhere (BBPOP 2012; Bull et al., 2013; IUCN 2016; Maron et al., 2025). Delivering on this promise requires that the ecological characteristics of the lost and gained biodiversity and their capacity to respond to management actions are accounted for in the design and evaluation of offset actions (Bull et al., 2013; Maron et al., 2012, 2025; Moilanen & Kotiaho, 2018). The goal of offsetting is usually No Net Loss (NNL) or net gain for biodiversity, where the biodiversity gains accrued from the offset actions fully match or exceed the losses (BBOP 2012; Bull et al., 2013, 2020; Moilanen & Kotiaho 2021). As offsetting typically trades immediate and certain losses with future and uncertain gains, reaching NNL requires larger offset areas compared to lost areas (Bull et al., 2017; Maron et al., 2016). In offset calculation, this is typically accounted for using offset multipliers (Maron et al. 2012; Moilanen & Kotiaho 2018; zu Ermgassen et al. 2021).

Offset multipliers are frequently criticized for lacking rigorous, quantitative evaluations of the biodiversity outcomes associated with offset actions (zu Ermgassen et al., 2019). Multipliers should be calibrated to achieve NNL, accounting explicitly for uncertainties, partiality, and time-lags associated with the expected biodiversity gains (Bull et al., 2013; Laitila et al., 2014; Maron et al., 2012, 2016). However, in practice, offsetting schemes account poorly for the differing biodiversity outcomes of the different offset actions and frequently use too small multipliers (Bull et al., 2017). In their review of the 108 offsetting schemes in different countries, Marshall et al. (2024) revealed that multipliers were mentioned only in 22 and specified in 11 policies. Multipliers ranged from 0 to 30, mostly between 0 and 6. Some offset policies specify risk of failure and time-lags in offset multipliers by, for example, discounting the future gains (Defra 2024), but many schemes do not specify these elements at all (Marshall et al., 2024). In addition to the offset action outcomes, offset multipliers can relate to for example conservation status of the impacted biodiversity, connectivity, probability of leakage, or location-based features (Moilanen & Kotiaho, 2018).

Offset actions include improving or creating the target biodiversity via restoration, maintaining or improving current biodiversity values via management, or by allowing passive recovery of already degraded habitats and avoiding future losses via protection (Maron et al., 2025). Here, we do not include avoided loss offsets (see Section 2.3.). Active restoration refers to actions that catalyse the development of the target ecosystem towards more natural state. Ecosystems can also recover passively after halting harmful human activity (Chazdon et al., 2024). The development after active restoration or passive recovery is often partial and never immediate (Curran et al., 2014; Elo et al., 2024; Prober et al., 2025; Spake et al., 2015), reaching the desired state can be hindered by, for example, failure in achieving favourable abiotic conditions to target biota, inadequate estimation of the (re-)dispersal capabilities of the desired flora and fauna, or various kinds of stochastic events (Elo et al., 2016; Halme et al., 2013; Maron et al., 2012). Ecosystems and their components typically respond in diverse ways to different restoration actions, or combinations of several actions (Haapalehto et al., 2014; Ramberg et al., 2023; Soomets et al., 2023; Spake et al., 2015). Similarly, different ecosystems require different management actions for preserving or increasing the current biodiversity values (Toivonen et al., 2015). To ensure balance between biodiversity losses and gains, offsetting schemes should acknowledge the variation in the outcomes across offset actions and habitats (whether habitat/vegetation types or species habitats) (Maron et al., 2012, 2016).

Moilanen & Kotiaho (2018) proposed using quantitatively defined biodiversity recovery over time as the basis for offset action specific multipliers, later formalized by Moilanen & Lehtinen (2025). This framework accounts for the effectiveness and time-lags of offset actions by modeling temporal responses of a biodiversity metric (e.g., habitat condition) after implementation of any given action (Moilanen & Lehtinen 2025) (Section 2.1). We call this framework the *Response-Based Habitat Hectare Assessment of Biodiversity Gains* (REHAB).

To operationalise the REHAB framework in practice, there are a few requirements to be fulfilled (Moilanen & Kotiaho, 2018). First, one needs a metric to assess biodiversity losses and gains. In principle, any relevant metric may be applied, but here we focus on an approach building on assessing habitat condition through ecological attributes important for biodiversity (Kangas et al., 2021; Moilanen & Lehtinen, 2025). Here, we refer to these types of metrics as habitat hectares (Parkes et al. 2003). Similar habitat or vegetation condition-based metrics are the most common evaluation criteria in offsetting schemes around the world (Borges-Matos et al., 2023; Marshall et al., 2024). The chosen condition metric should be relevant for the ecology of the target biodiversity and enable verifying and monitoring offset action outcomes in the field (Contos et al., 2025; Marshall et al., 2024; Moilanen et al., 2024; Strange et al., 2024). Another requirement is knowledge about the ecological responses that describe how the chosen condition metric evolves through time after offset actions have been carried out. The use of responses has been scarce in offsetting due to the practical challenge of acquiring realistic and justified response information (but see e.g. Mokany et al., 2025 for ecosystem accounting). Ideally, one would use empirical field data from systematic monitoring of offset actions (Moilanen et al., 2024) and meta-analyses thereof (Prober et al., 2025). However, one must often rely on other sources of evidence and expert elicitation (Hemming et al., 2018; Martin et al., 2012).

In this paper, we present a practical approach for deriving response functions, consequent offset multipliers, and ecological attributes and their condition, to operationalise the REHAB framework within offset policy. We begin with a detailed outline of the framework, followed by a description of how ecological attributes, their condition matrices, and response data were compiled through large-scale expert elicitation. The approach was developed during the implementation of Finland’s biodiversity offsetting policy and provides outcome estimates and consequent offset multipliers for all Finnish habitat types and major restoration and management practices. Although our examples are from typical boreal and hemi-boreal ecosystems, the approach presented here can be applied globally to any biodiversity (species’ habitats, biotopes and ecosystems).

## 2 Materials and methods

### 2.1 The REHAB framework

Offset multipliers define the ratio between the magnitude of biodiversity loss at the impacted area and the area required to deliver equivalent biodiversity gains with any given offset action (Moilanen & Lehtinen, 2025). In the REHAB framework, units of losses are habitat hectares (hha) that are defined as the product of area (ha) and ecological condition (hha/ha). Ecological condition refers to the status of those ecological attributes that are central for supporting biodiversity in each habitat. Mathematically, condition is expressed as a number between 0 and 1 (Moilanen & Lehtinen, 2025). Thus, the condition of one hectare of habitat, of which the ecological attributes important for biodiversity have been degraded by half (0.5), would equal to 1 ha × 0.5 hha/ha = 0.5 hha. Different ecological attribute values can be aggregated into a condition value in various ways (Borges-Matos et al. 2023).

Quantifying offset gains requires an estimate of how any given offset action impacts the ecological condition of any given habitat over time. This is called a *response*, a function specific to each action–habitat pair (Figure 1a). From the response, the *average gain* [hha/ha] over time is calculated as the arithmetic mean of the cumulative improvement (Figure 1b) for a given evaluation time frame (e.g., 30 years). The evaluation time frame is needed to assess the equivalence between gradually accumulating offset gains and usually immediate losses (Moilanen & Kotiaho, 2018). Note that the length of the time frame is ultimately a societal subjective decision that needs to be defined in the underlying offset policy. The shorter the evaluation time frame is, the larger the resulting multipliers become for reaching NNL, while too long evaluation time frame increase the uncertainty around achieved gain and decreases the credibility of the offset scheme (Moilanen & Lehtinen 2025).

**Figure 1.**
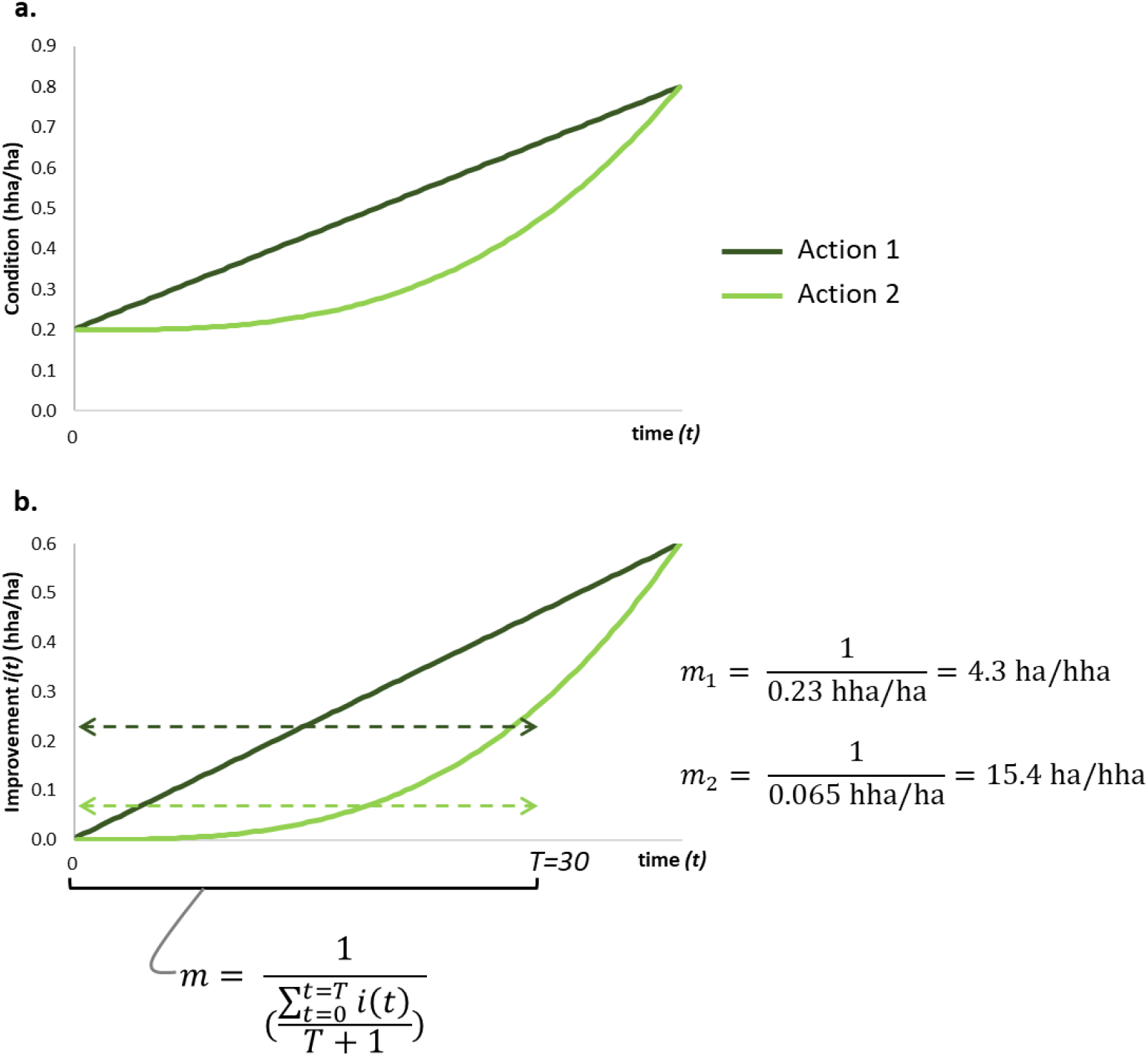
An illustration of the REHAB approach for calculating offset action multipliers (*m*), originally described by Moilanen & Lehtinen (2025). (**a**) Two alternative actions can be carried out to improve a habitat’s ecological condition (hha/ha) at a given site. Both actions improve the condition the same amount, from 0.2 to 0.8 hha/ha, after equal number of years (*t*) but the habitat condition’s response to action 1 (dark green) is linear, i.e., the condition improvement is constant over time, and covex to action 2 (light green), i.e., response is initially slow but accelerates through time. (**b**) From the response, we derive the cumulative improvement in habitat condition *i(t*) (hha/ha) through time. This is used to calculate the the average gain, i.e., the mean cumulative condition improvement over the evaluation time frame *T*, marked by dashed horisontal lines. Offset action multiplier *m* (ha/hha) is then calculated as 1 divided by the average gain. Calculations for the action-specific multipliers *m*_*1*_ and *m*_*2*_ are shown on the right, using an illustrative evaluation time frame of 30 years. Action 1 results in greater average improvement and thus smaller offset multiplier (6.7) compared to action 2 (55.6).

Using the average gain from the response, one can then calculate the offset action multiplier as the inverse of the average gain to define the area needed to reach NNL with the action (Moilanen & Lehtinen, 2025) (Figure 1). For example, if the offset action’s average gain is 0.32 hha/ha over a given evaluation time frame, the respective multiplier needed to produce NNL is 1 / (0.32 hha/ha) = 3.125 ha/hha. Hence, offsetting a loss of 2 hha with this offset action requires that the action is implemented on 3.125 ha/hha × 2 hha = 6.25 ha to reach NNL (ignoring other possible offset multipliers).

### 2.2. Biodiversity offsetting in Finland

Finland introduced voluntary biodiversity offsetting in its legislation for the first time during the revision of the Finnish Nature Conservation Act in 2023 (Finlex 9/2023). The legislation requires that biodiversity losses and gains are measured in habitat hectares as described above, and that the offset gains are estimated based on the response to the offset actions (Decree of the Ministry of the Environment on Voluntary Ecological Compensation; Finlex 933/2023). The Decree sets the evaluation time frame for calculating the average gain to 30 years and after a site has been used as an offset, it will be protected in-perpetuity. The legislation directly creates a need for i) defining appropriate ecological attributes for each habitat and ii) deriving response estimates for each of the habitats to relevant offset actions. Currently, Finnish legislation does not recognize avoided loss offsets. However, REHAB framework can be applied also to quantify gains from avoided losses (Moilanen & Lehtinen 2025).

### 2.3. Applying REHAB for Finnish habitat types: broad overview

Our aim was to apply the REHAB framework for Finnish habitat types, in support of operationalizing biodiversity offsetting in Finland. As most Finnish offsetting cases are likely to target habitat types (Suvantola et al. 2024), we aimed to produce response estimates for all relevant active restoration and management actions as well as passive recovery in all broad ecosystem groups.

There are 388 natural habitat types listed in the current version of the Finnish Red List of Habitat Types (Kontula & Raunio 2019), grouped into eight broad ecosystem groups: the Baltic Sea (marine); the Baltic Sea coast; inland waters and shores; mires; forests; rock outcrops and scree (hereafter rocks); seminatural grasslands and wooded pastures (hereafter grasslands); and fell habitats. As part of the process of producing ecological attributes, their condition matrices, and response estimates for offsetting, we further divided the inland waters into lakes and ponds; springs; streams; and shores. We excluded pelagic habitats, sea ice, or waterfalls because, based on expert judgement, offsetting is impossible in these habitats.

We used large-scale expert elicitation to define the most suitable ecological attributes and to build condition assessment matrices and action responses for the Finnish habitat types. The large number of experts served the purpose of ensuring both the quality of the work and its legitimacy among Finnish practitioners. In total, 111 ecology, restoration and habitat type experts participated in the collaborative effort, including scientists and experts from Finnish state research institutes, state and municipal environmental officers, consultant companies, and NGOs. During the process, experts worked in thematic small groups focusing on the different broad ecosystem groups listed above. The number of experts in one group ranged from 5 (rocks) to 25 (forests).

The expert engagement was carried out in workshops from November 2022 till September 2023. The first workshops for both condition metrics and responses were in-person events, followed by online workshops, and additional discussions via email. The workshops included a brief introduction to offset calculations and offsetting in general. In total, 22 workshops were held. Experts were also asked to comment on the draft versions of the materials produced throughout the process.

### 2.4. Condition metrics

#### 2.4.1. Scope of condition metrics

For each habitat type, we aimed to create an assessment matrix, where individual ecological attributes are scored on the scale of 0.1–1; 1 meaning pristine and 0.1 nearly collapsed condition. The total condition of a habitat type patch is then the weighted average of individual ecological attribute scores. The condition assessment matrices were meant for practical field work. They should be accurate enough for condition assessments to produce ecologically credible results, yet not too laborious to be operational in practice.

#### 2.4.2. Expert engagement

The experts compiled the condition assessment matrices by defining the following information for each habitat type: (i) which ecological attributes are required to assess the ecological condition of a site, (ii) how each attribute is divided into condition classes on the scale of 0.1–1, (iii) verbal descriptions for different condition classes (2–5 classes per attribute; minimum requirement was to describe condition classes 0.1 and 1), and (iv) relative weights of ecological attributes, based on their ecological importance, which are used for calculating the final condition score. For simplicity, only relative weights 2 and 1 were given as an option.

The iterative process of formulating the condition matrices was as follows: First, a draft proposal was produced by researchers, which mainly relied on existing evaluation frameworks such as the assessment criteria for the EU Habitat Directive’s habitat types (Airaksinen & Karttunen 2001) and the Finnish forest assessment criteria used in conservation planning (Syrjänen et al. 2016). The proposal was then modified or remade by experts in workshops. During this process, experts were given the option to merge assessment matrices for ecologically similar habitat types within the same broad ecosystem group. After this, researchers harmonised the verbal descriptions of the assessment matrices which were again reviewed by the experts. The assessment matrices were published as a draft for public scrutiny in April 2023. Based on the received feedback, researchers finalised the matrices and wrote a detailed interpretation guide. Experts were again given the chance to review the guide before it was published.

### 2.5. Responses

#### 2.5.1. Parameters needed for response functions

For restoration and passive recovery, the offset gain equals the improvement in the focal habitat type’s condition through time (Figure 2a, b). They are thus cumulative functions that start from 0 at the time of completing the action and gradually increase into some response-specific maximum value (Moilanen & Lehtinen, 2025).

**Figure 2.**
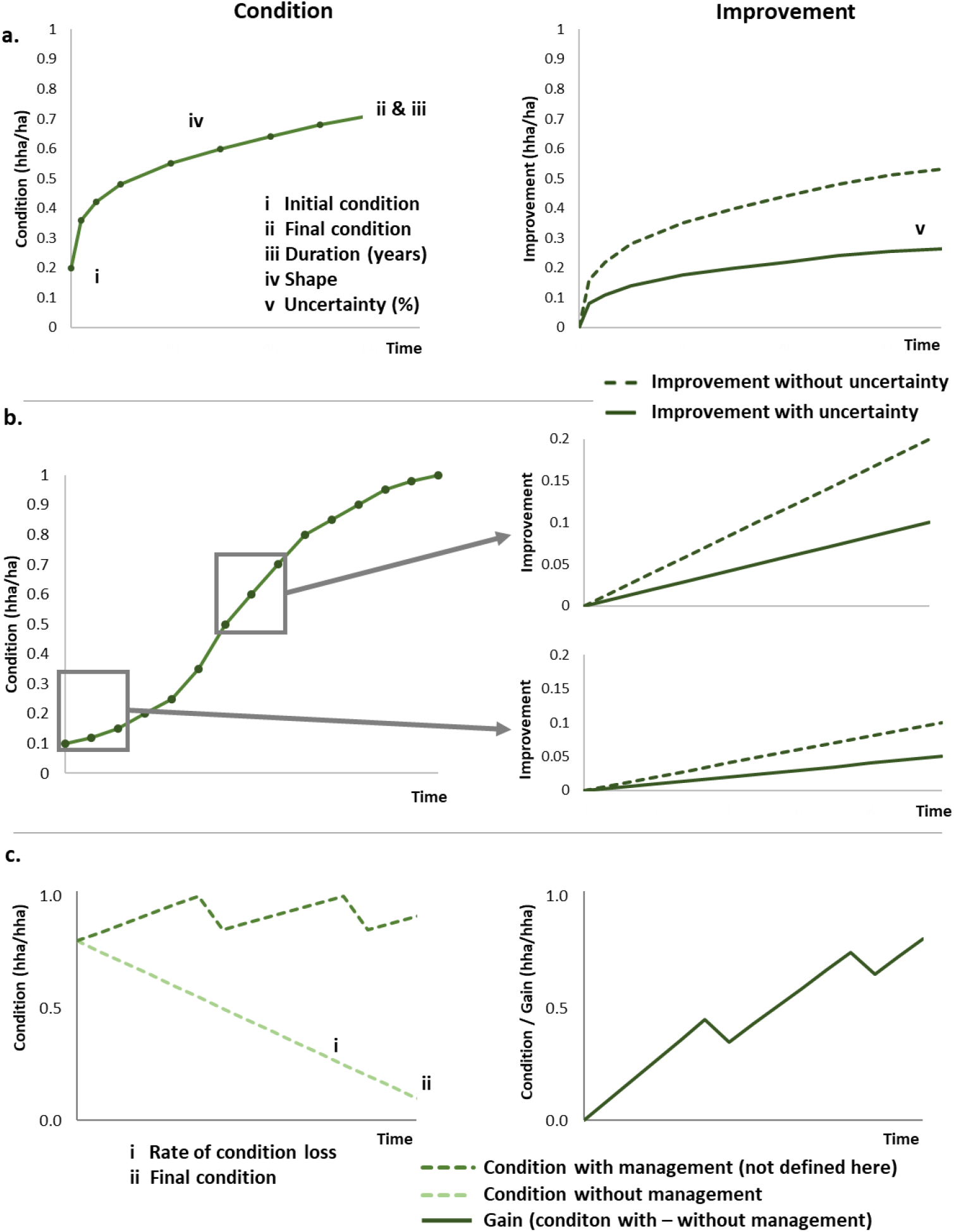
(**a**) Calculation of offset gains from restoration actions. One must define the initial condition (i), the final condition (ii), the duration between end points i & ii (iii), and shape of the function (iv). From this function describing condition development through time, the cumulative improvement is calculated (right panel). Finally, uncertainty (v) decreases the gain that is accounted from the condition improvement from the restoration action. See Supplementary 1 for mathematical formulations. (**b**) Calculation of offset gains for passive recovery, of which the improvement depends on the initial condition. First, one needs an estimate for how the condition develops via natural succession, which can be compiled from parameters i– iv presented in (a). Response function is then calculated as the improvement from the initial condition onwards (here, 0.1 and 0.5 hha/ha; right panels). Uncertainty decreases the expected gain from that estimated future improvement. (**c**) A schematic illustration of a response function for management. First, one must define the baseline function, i.e., how fast the target habitat type’s condition degrades without management (i), and the final condition habitat type ends up (ii). Then, a response function with management is needed. The gain is the difference between these two functions. Here, only baseline functions without management were defined.

We used the following basic parameters to guide the building of response functions for restoration and recovery (Figure 2a):

i. *Minimum initial condition*. The lowest condition, where the action is still appropriate for the habitat type.
ii. *Final condition*: the maximum condition that the target habitat type can eventually reach when the action is conducted to a habitat in the lowest possible initial condition (i).
iii. *Duration*: the number of years the target habitat type takes to develop from the minimum initial (i) to the final condition (ii).
iv. *Shape* of the response function. We defined six simple but ecologically justifiable alternatives: linear, convex (steeper and less steep), concave (steeper and less steep), and sigmoid (Figure 2b, Supplementary 1).
v. *Uncertainty*: the risk (%) that the trajectory defined with parameters i–iv is not reached with the present level of ecological knowledge and practical expertise regarding the offset action and the habitat type.

The uncertainty value is used to reduce the expected offset gain from each action: e.g., if uncertainty is 30%, only 70% of the estimated condition improvement for the said habitat type is counted as gain each year (Supplementary 1).

#### TOPIC: parameters needed for management responses

In contrast to active restoration, management is a repetitive action, without which the habitat type’s condition would decrease (Moilanen & Lehtinen 2025). A management offset gain is defined as the difference in two responses: response of habitat condition to management, and a baseline response when left without management (Figure 2c). For simplicity, we assumed a linear decrease in condition until a habitat-specific minimum condition is reached, which allows defining the yearly rate of condition loss (Supplementary 1). For that, the following parameters are needed:

i. *Final condition*: the condition class the target habitat type eventually ends up when left without management.
ii. *Duration*: the number of years it takes to reach the final condition (i) when a site in perfect condition (1 hha/ha) is left without management.

Note that the parameters i–ii only define the baseline responses that would result if the management was ceased. The responses to management, i.e. how the condition improves or is maintained in time due to management actions, depend on the case-specific management regime and intensity and need to be defined case-by-case. Uncertainty associated to management gain is accounted for when defining responses to management. We therefore did not include uncertainty here in the baseline response estimates to avoid counting the uncertainty twice.

#### 2.5.2. Expert elicitation for response parameters

First, the experts identified a list of relevant offset actions for different habitat types in workshops. The response parameters per action–habitat type pairs were then defined in a combination of a questionnaire and workshops.

In the questionnaire, experts individually went through each identified action and i) assessed which habitat types it applies to, and ii) estimated its response parameters (Section 2.5.1) for each habitat type, or a group of habitat types. To do this, experts were given some practical assumptions regarding the responses. First, it was assumed that the given action is always adjusted to the local circumstances and executed perfectly. The experts were then asked to assess the average trajectory of the focal habitat type. It was thus acknowledged that offset actions improve the condition more and faster in some places and less and slower in others, even when perfectly executed. Estimates for the uncertainty include, in part, how well-known or difficult the action is to execute. We asked the experts to specify the duration of the response separately for southern and northern Finland for those habitat types that are distributed across the country, because the climatic conditions differ between the regions (Supplementary 2).

In the following workshops, experts were presented with the anonymised answers of the questionnaire and asked to discuss and develop them further. The lowest and final condition and the shape of each response were discussed in groups until a consensus among experts was reached. Following the precautionary principle, experts were advised to choose the more conservative option in uncertain situations.

Duration and uncertainty were defined following the IDEA protocol for structured expert elicitation (Hemming et al., 2018). In the questionnaire, the experts gave their upper, lower and best estimate for duration and uncertainty, and their confidence rate for duration. In the workshops, again the anonymised answers were presented (rescaled to 80% confidence interval to ensure comparability; Hemming et al. 2018) and discussed. After discussion, each expert was asked to give their new best estimate for duration and uncertainty. The final values were the averages of the workshop answers (Hemming et al., 2018). When the response’s average duration estimate exceeded 10 years, the average was rounded to the closest 5 years. Similarly, when average uncertainty estimate exceeded 10%, it was rounded to the closest 5%. This was done to avoid pseudo accuracy.

The response parameters were in some cases later discussed and revised by the thematic groups via email. Afterwards, the researchers wrote a verbal description and justification for each response estimate, which the experts were given the chance to review before making the estimates public.

### 2.6. Calculation of offset action multipliers

Finally, we calculated the offset action multipliers for each response estimate. The multipliers were based on average condition improvements calculated over the first 30 years from executing the action, as defined in the Finnish offset legislation (Decree of the Ministry of the Environment on Voluntary Ecological Compensation 4 §).

We note that offset actions may be carried out at sites that are in better ecological condition than the minimum condition defined by the experts. This begs the question of how the responses should be applied to different starting conditions. Based on expert views, we chose to apply the responses and calculate the corresponding offset action multipliers differently for active restoration and passive recovery. In active restoration, local abiotic and/or biotic conditions are modified to create a change in some or all the ecological attributes, thus facilitating a change that would not occur without restoration. Although the development is likely faster or more certain in intermediately degraded than severely degraded sites, we expect the condition improvement to follow a somewhat same trend regardless of the initial condition. We therefore assumed that for restoration, the same response would apply regardless of the initial condition of the offset site (Figure 2a). This assumption however applies only if one uses conservative response estimates that are not overly optimistic in degraded sites. Importantly, when the action is implemented at sites in higher initial conditions, the response values may need to be capped to 1 before calculating the improvement and average gain (Supplementary 5). Further, not all actions can be applied to higher conditions: e.g., establishing new ponds can only be applied to sites that do not have an existing pond, i.e. to initial condition of 0. For simplicity, we report active restoration multipliers in the main text using only the lowest initial condition. For illustration, we show multipliers for all starting conditions in Supplementary 5.

With passive recovery, the average condition improvement depends on the initial condition. In passive recovery, the target habitat type recovers towards natural state via natural succession. In habitat types that can recover by themselves, this development starts immediately and depends on the ecological characteristics already present on the site. Using forests as an example, if a clearcut is set aside, it takes decades to centuries before ecological attributes considered important for biodiversity (large deadwood, old trees, etc.) develop. Setting aside a mature forest would result in these features being present much faster. To account for the effect of starting condition, we therefore used the response defined by experts for the lowest possible condition, find the time point that matches the starting condition of the site and calculate the expected condition improvement from that point onwards (Figure 2b). From this future improvement, uncertainty is reduced, and average gain is calculated. Thus, for passive recovery, several offset multipliers are defined per habitat type.

For comparison, we calculated offset action multipliers for management actions assuming that the target areas would be constantly managed to maintain their perfect condition (1 hha/ha) and excluding uncertainty. Thus, the multipliers presented here are the theoretical minimums and, in reality, multipliers for management are likely to be larger than the ones presented here.

## 3 Results

The original condition metrics and response materials are publicly available in Finnish in Zenodo (https://zenodo.org/records/14409001).

### 3.1. Condition metrics

There were total of 37 condition assessment matrices defined (Supplementary 3). One assessment matrix can be applied to several ecologically similar habitat types. Within broad ecosystem categories, number of matrices ranged from one (springs, streams, shores) to nine (rocks). Number of ecological attributes within a condition assessment matrix ranged from one (scree) to eight (moist herb-rich forests), mostly between four and seven. Table 1 provides two examples of the matrices.

**Table 1.**
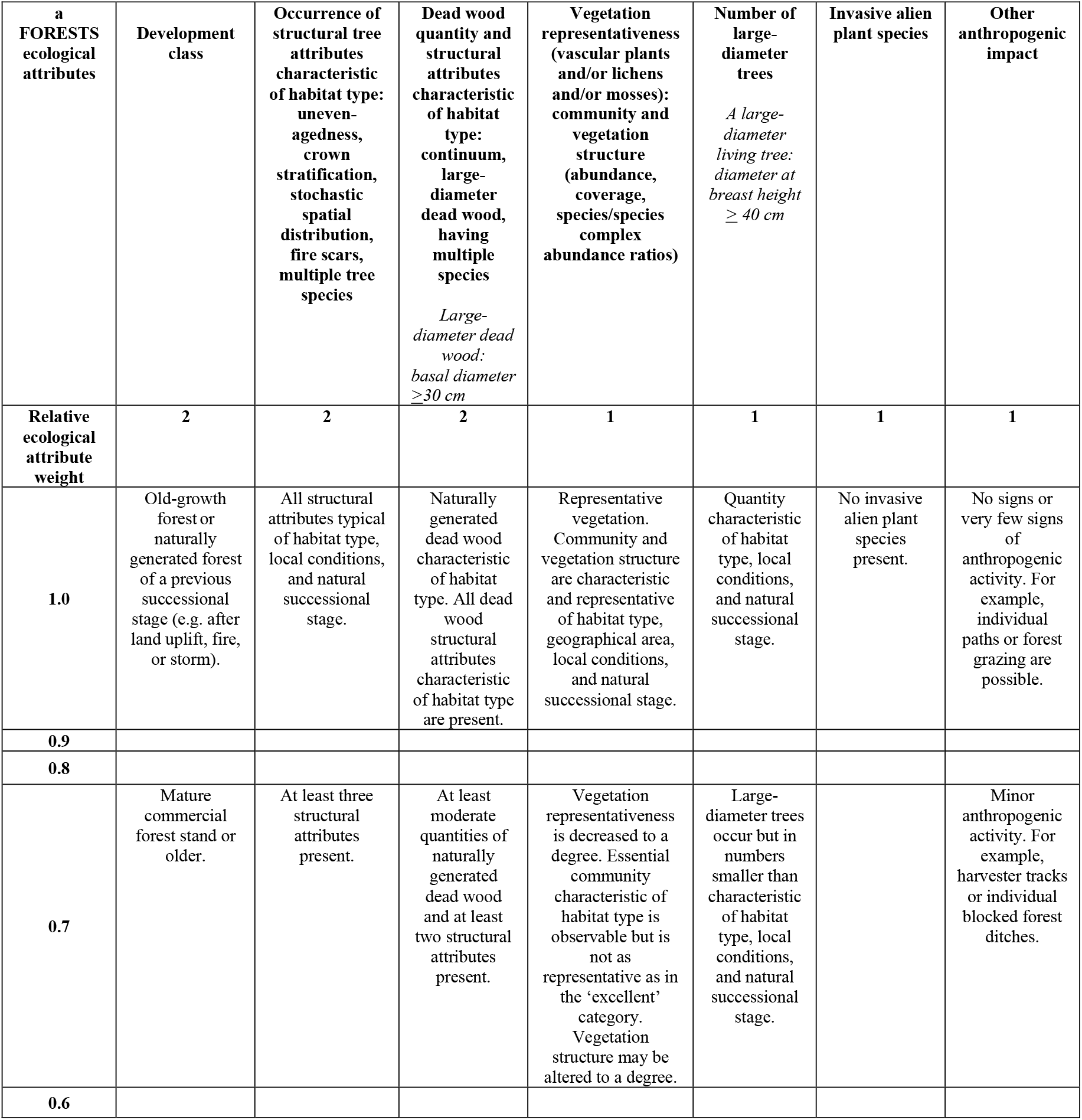
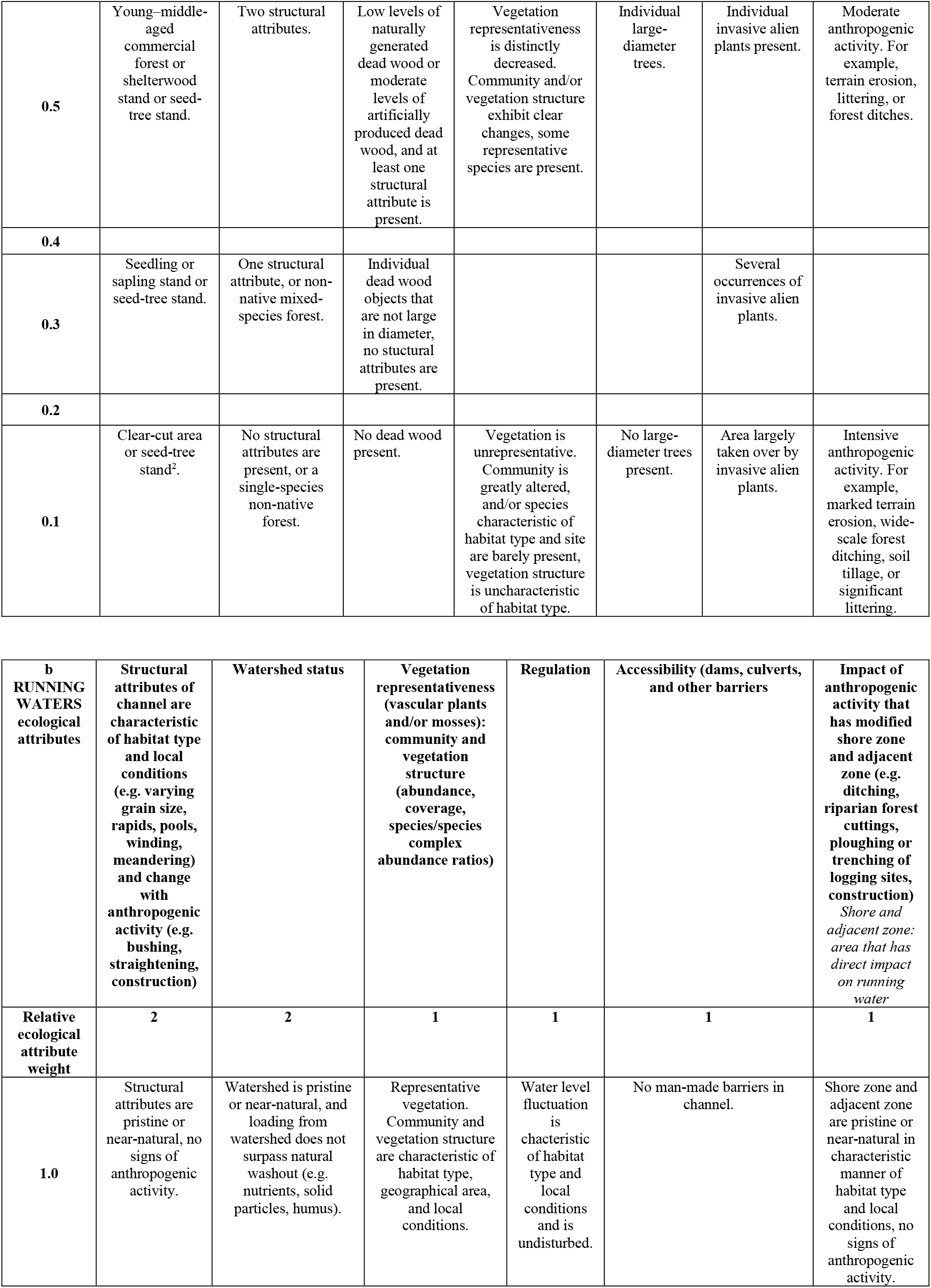
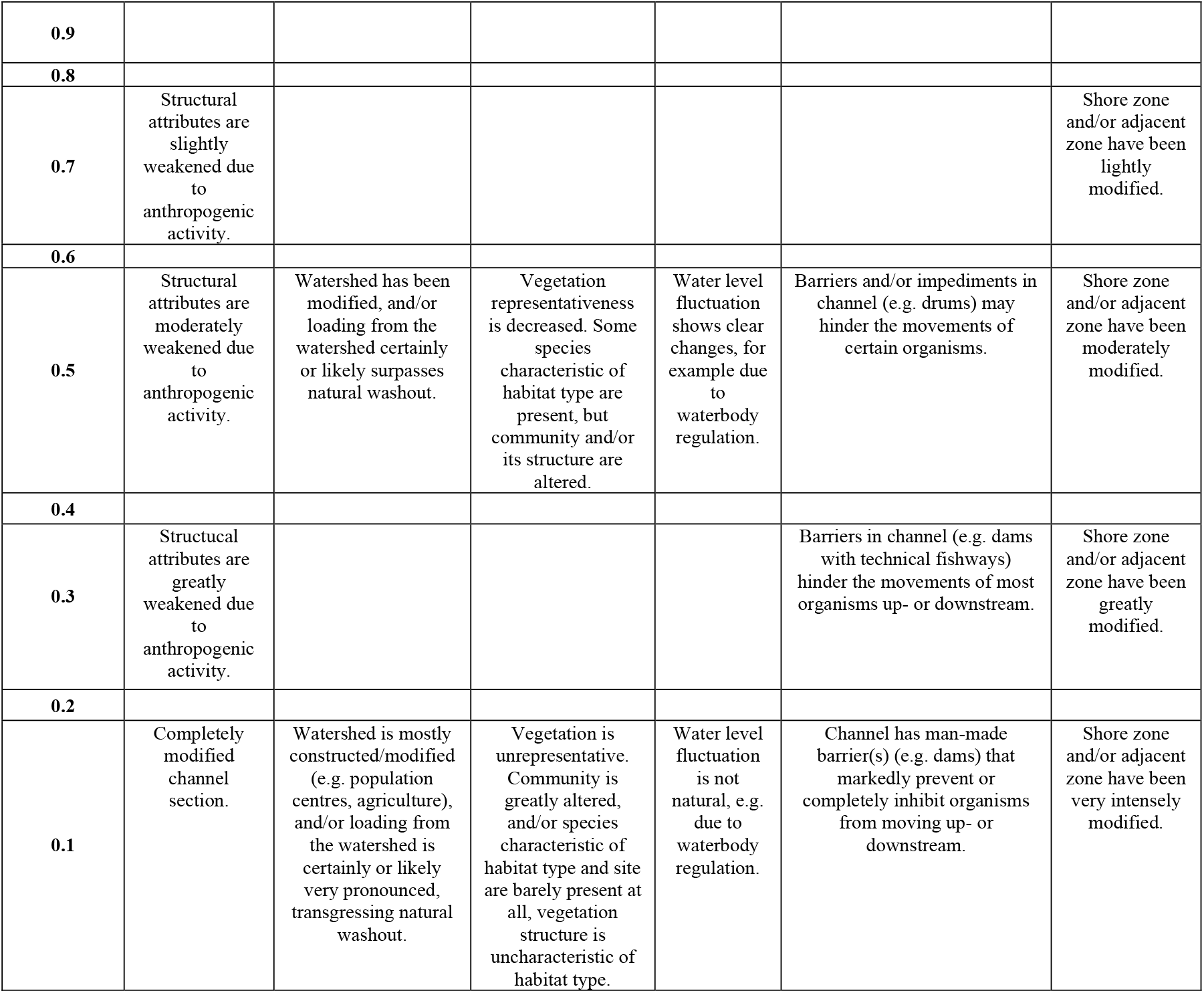
Examples of condition assessment matrices for (a) forests and (b) running waters (see Supplementary 3 for all matrices).

Most of the ecological attributes can be assessed on the field, but some require GIS assessments (e.g., lakes’ catchment areas). Vegetation representativeness was a common attribute, applied to nearly all habitat types, in which vegetation is assessed against the pristine vegetation that is described for each habitat type in the Finnish Red List of Habitat Types.

Of note, the metrics defined here were adopted in the Finnish legislation as mandatory condition assessment criteria for habitat types (Decree of the Ministry of the Environment on Voluntary Ecological Compensation Annex I; Suvantola et al. 2024).

### 3.2. Responses and multipliers

In total, 216 responses for action–habitat type pairs were compiled (Supplementary 4). The number of actions per ecosystem group ranged from two (fells) to seven (forests).

Restoration actions (182) were deemed appropriate for all habitat types except for fells. Passive recovery (19) was seen as possible in forests, primary forests in land uplift coasts, and fells. Management actions (15) were identified appropriate for grasslands and some forest types. Furthermore, baseline responses for preserving forests that shade rock faces were defined during the process. Although those responses relate to avoided losses, here they were defined like management, as losing the shade would lead to a baseline loss in condition due to microclimatic changes.

The critical components of the action responses are the total improvement, duration, shape and uncertainty of the response, which also influence the final offset action multiplier. These components all varied across the restoration and recovery responses compiled here (Figure 3; Supplementary 4). The total improvement ranged from 0.11 (removal of small culverts in streams) to 1 hha/ha (e.g., establishing new flooded mires). These varied relatively little among actions in inland water habitat types and rocks, and greatly in the Baltic Sea coast, mires, and grasslands (Figure 3a).

**Figure 3.**
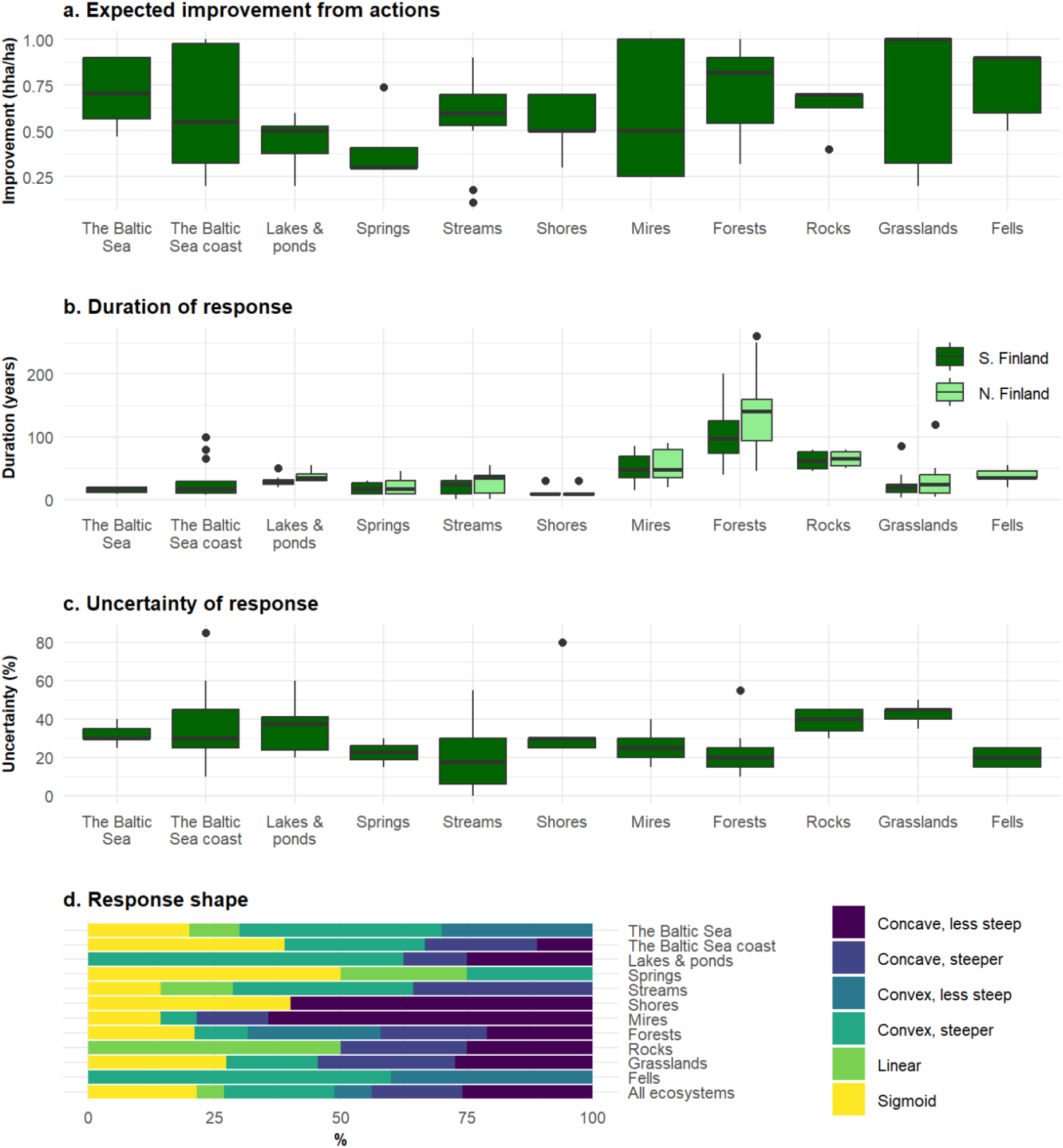
Distributions of parameters of the restoration and passive recovery response functions, grouped per ecosystem category. (**a**) Total improvement expected from action, calculated as final condition minus lowest possible initial condition. (**b**) Duration of response, shown as years to reach the final condition from the lowest possible initial condition. Duration is defined separately for southern and northern Finland while other parameters were considered the same between the regions. (**c**) Uncertainty of response (%). (**d**) Proportions of different shapes in the response functions.

Duration of restoration and recovery actions varied among ecosystem categories being longest in forests, mires, and rocks (Figure 3b). The duration ranged mostly between 15–65 years. The fastest action was the removal of small culverts in streams, which has an immediate effect (duration 1 year to maximum improvement) and slowest in passive recovery of North-Finnish barren heathland forests, which were estimated to take 260 years to reach the natural condition from a clearcut.

Uncertainty in achieving the assumed improvement varied mostly between 20 and 40% between actions (Figure 3c). The uncertainty varied the least in grasslands and the most in the Baltic Sea coast and streams. Some actions were assigned with low or no uncertainty, the extreme being the removal of small culverts in streams (0% uncertainty). On the other hand, some actions were deemed very poorly known and extremely uncertain, such as establishing new coastal tall-herb meadows (85% uncertainty).

Across all habitat types, 88 (44%) responses were estimated to follow concave and 59 (29%) convex shapes, either steeper or less steep versions (Figure 3d). Sigmoid shape was deemed the most appropriate for 43 responses (21%) and linear shape to 11 responses (5%).

From the responses, a total of 346 offset action multipliers were calculated, including one multiplier for each restoration and several for each passive recovery response. There was substantial variation in the sizes of offset multipliers for restoration actions (Figure 4a). Most varied between 2 and 40 ha/hha, but for three responses they exceeded 100 ha/hha (restoration of eutrophicated lakes in southern and northern Finland, and restoration of coastal wooded dunes; 149.3, 198.8, and 191.2 ha/hha, respectively). Within ecosystem groups, variation in the multiplier values was the smallest for the Baltic Sea habitat types, (1.8–5.3 ha/hha) and the greatest for lakes and ponds in northern Finland (2.8–198.8 ha/hha). The results are based on responses starting from the lowest possible initial condition. The general patterns of the multipliers were the same regardless of initial condition (Supplementary 5).

**Figure 4.**
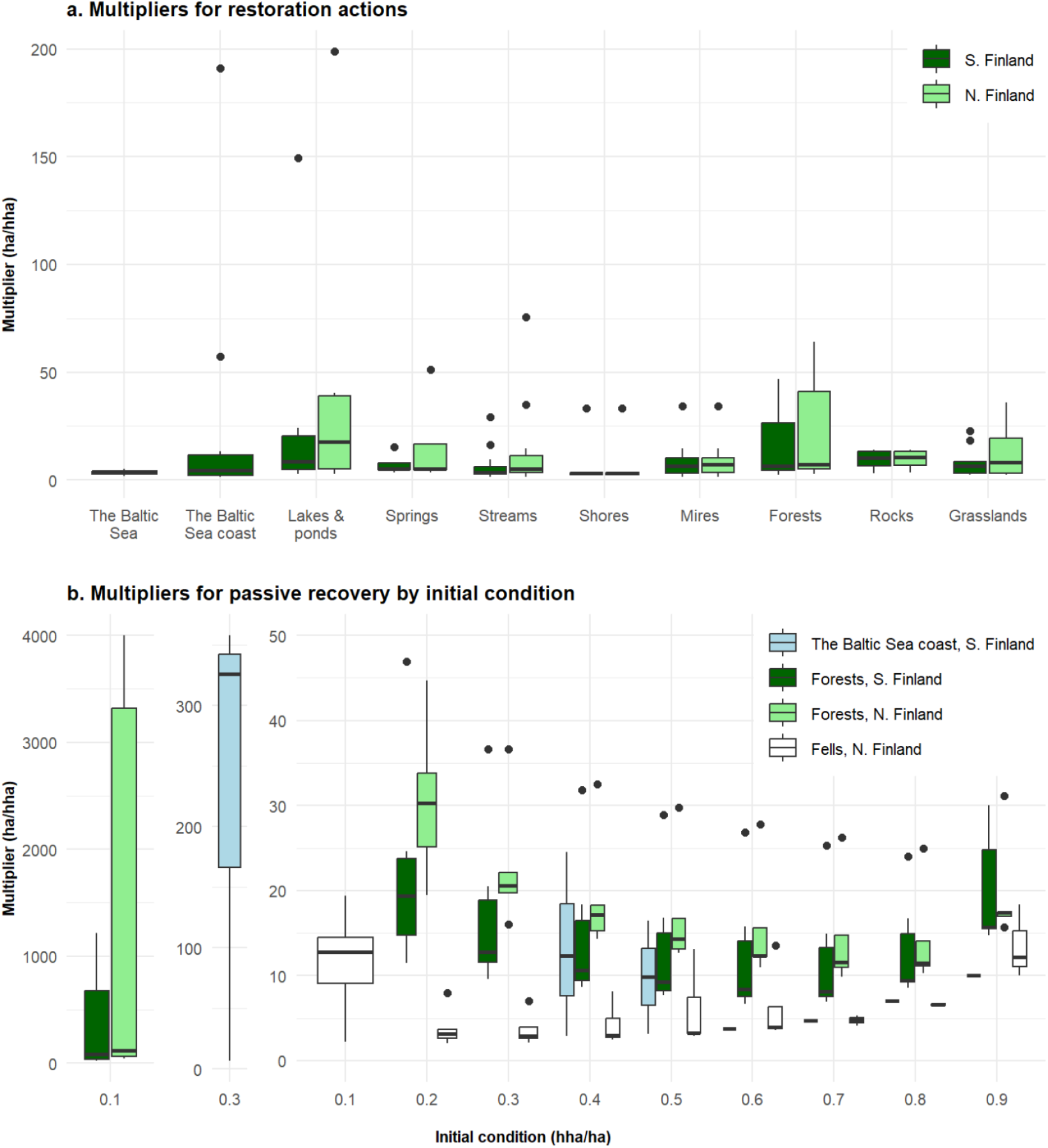
Distribution of offset action multipliers for (**a**) restoration and (**b**) passive recovery actions, grouped per ecosystem group and region (southern/northern Finland). In (**b**), passive recovery offset action multipliers depend on the initial condition of the target habitat type. Multipliers for forest types from the initial condition 0.1 and the Baltic Sea coast habitat types from the initial condition 0.3 are plotted separately, as their values are on a very different scale compared to other cases.

Multipliers for passive recovery varied even more, between 2.1– 3,999.8 ha/hha (Figure 4b). Multipliers generally ranged between around 8–30 ha/hha for southern Finnish forests, around 10–50 for northern Finnish forests, around 3–25 for coastal forests, and around 3–30 for fells. However, when the initial condition was low, the multipliers for forests could be vastly greater: multipliers ranged from 6.9 (coastal forest types in the lowest possible initial condition of 0.3 hha/ha) to 3,999.8 ha/hha (barren heath forests in the condition of 0.1 hha/ha). Compared to these extremes, the multipliers decreased rapidly for forests in higher initial condition.

Figure 5 shows the response parameters and multipliers for management actions. Without management, the condition of grasslands decreases the most (Figure 5b); the loss was estimated to be 0.8 hha/ha for dry meadows and complete (1 hha/ha) for moist meadows. For forests, the estimated loss was 0.3–0.7 hha/ha, and 0.7 hha/ha for rocks. For forests and grasslands, this baseline loss in the absence of management was estimated to take decades (Figure 5c), whereas rock faces were estimated to lose their condition in two years after losing shade. Multipliers ranged from 1.5 (preservation of shading forest in rock faces) to 14.4 ha/hha (removal of spruce *Picea abies* in broadleaved deciduous herb-rich forests).

**Figure 5.**
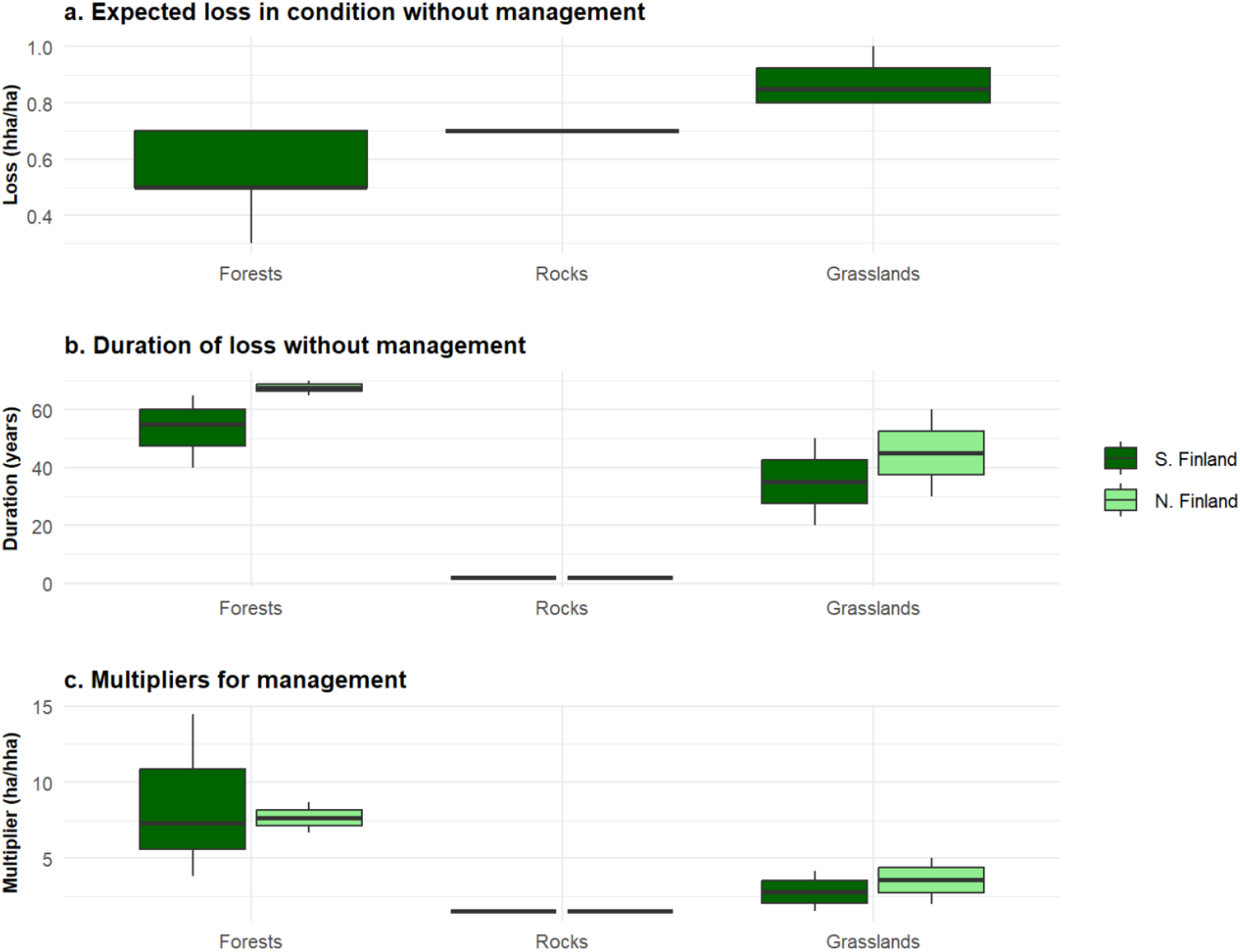
(**a**) Expected loss in ecological condition (hha/ha) that each habitat type face without management. (**b**) Duration of the loss without management, expressed as years to reach the final condition from the natural condition in the absence of management, grouped per ecosystem category and region (southern/northern Finland). (**c**) Distribution of offset action multipliers for management, grouped per ecosystem category and region (southern/northern Finland). These multipliers represent the theoretical minimums as they rely on an assumption that each habitat is maintained constantly in a natural condition with management and exclude uncertainty.

## 4. Discussion

Building on the earlier work of Moilanen & Kotiaho (2018) and Moilanen & Lehtinen (2025), we present an operationalization of the framework we call the ‘Response-Based Habitat Hectare Assessment for Biodiversity Gains’ (REHAB). We define response estimates and respective offset action multipliers for 216 action–habitat type pairs to support an emerging national biodiversity offsetting scheme in Finland. Importantly, we demonstrate that the REHAB approach, although seemingly complex at first glance, can be applied in practice to support ecologically informed offsetting and, more generally, restoration and management planning. Furthermore, as our work has been directly adopted into practice and legislation in Finland, it serves as an example of a successful science–policy implementation.

Our results show that there is no one-size-fits-all multiplier that can capture the ecological realism of biodiversity offsetting (Figure 3). For instance, the multiplier 6, a common upper multiplier identified by (Marshall et al. 2024), would overshoot offsetting needs in 36% of cases, being unfair for practitioners, and undershoot in 58% of the cases, being unfair for nature. Offset multipliers also varied greatly between actions, not only between habitat types (Figure 2). Thus, applying fixed multipliers within habitat types regardless of the specific offset action, such as in the Biodiversity Net Gain tool (Defra 2024), is prone to the same lack of ecological realism.

It is also clear that the multipliers currently used in offset policies are too small to deliver NNL. Even the multiplier 30, sometimes mentioned as the highest one used (Bull et al., 2017; Marshall et al., 2024), would be too small in 13% of the Finnish cases. We note that here we focused solely on the offset action multiplier and excluded others, such as threat, measurement uncertainty, or partial additionality (Moilanen & Kotiaho 2028), which would further increase the final offset multipliers required for NNL. We also did not time discount the response functions to penalize biodiversity gains achieved in the future (Laitila et al. 2014; Moilanen & Lehtinen 2025) as it is not required in the Finnish offsetting legislation. Using time-discounting would further increase the offset action multipliers.

The offset action multiplier depends on the effectiveness (total condition improvement achieved), duration, shape, and uncertainty of the action for the target habitat type (Figure 2). Poor performance in any of these parameters increases the offset action multiplier. For instance, the ecological impact of removing culverts in small streams is almost immediate (duration 1 year) and completely certain (uncertainty 0%), but it increases the stream habitat type’s condition only by 0.11 hha/ha. Thus, the consequent multiplier is relatively high, 9.4 ha/hha. Uncertainty values relate here not only to the direct risk of failure but also to the current level of knowledge among practitioners. Extremely uncertain actions are unattractive until known better, thus hopefully incentivizing research and monitoring of those actions (Moilanen et al., 2024). Worth noting is that linear response was seen appropriate for only about 5% of the response estimates (Figure 4d). Although linear response can be a practical starting point for response estimates, based on our expert solicitation it is unlikely to very often appropriately describe the ecological changes over time.

The REHAB framework is essentially about accounting for the ecological effectiveness and efficiency of the intended actions in offset planning. For instance, in eleven cases (3%), the offset action multiplier was >100 ha/hha. The largest multiplier was almost 4,000 ha/hha for passive recovery of northern Finnish barren heath forests in totally degraded condition (0.1 hha/ha). Thus, a loss of one hha would require setting aside 40 km^2^ of degraded barren heath forests. Given the specific action is so inefficient, it will likely not be used as an offset at all.

One key strength of the REHAB framework is that it brings transparency to offsetting and can limit the use of inefficient offset actions. However, it is crucial to acknowledge that not all offset actions are applicable in all cases. Many actions have ecological prerequisites for achieving the desired development (e.g., planting underwater meadows will not succeed if the water quality is too poor). Similarly, a site might be in such good condition that it will not benefit from restoration (Chazdon et al., 2024). Higher initial condition may also increase restoration multipliers because the total condition improvement is lower compared to sites in low initial conditions (Supplementary 5). However, many restoration actions likely have an upper limit after which the said action becomes ecologically inappropriate. All offset actions should primarily be chosen based on local considerations about what is feasible and what truly benefits the local biodiversity, if offsetting is really meant to mitigate the biodiversity crisis.

The obvious limitation of the REHAB framework is the extensive need for ecological information and the more complicated offset calculations compared to fixed multipliers. The first practical challenge is to acquire the response functions. However, as our work demonstrates, the required information can be compiled relatively feasibly. We relied largely on expert engagement due to lack of empirical ecological data, as we would expect to be the case in all other countries as well. Expert elicitation is a useful source of information in conservation planning, and methods exist to overcome its inherent biases (Hemming et al., 2018; Martin et al., 2012). In our case, the IDEA protocol proved useful in acquiring information and increasing shared knowledge and agreement between experts in a relatively short time. Furthermore, simulations, meta-analyses and empirical data accumulating from restoration monitoring schemes (e.g., Spake et al. 2015; Shorohova et al. 2024; Elo et al. 2024) can be used to adjust response estimates over time. National offset monitoring schemes (Moilanen et al., 2024) would also greatly benefit offset planning in the future.

There are also ways to refine our responses estimates. They could be assigned per ecological attribute instead of the condition as a whole, which would add ecological details and make on-the-ground evaluation easier in the future (Maseyk et al., 2016). Responses can also be made more detailed to account for, for example, how local conditions affect the effectiveness or uncertainty of actions (Mokany et al., 2025). Yet, offset policies must balance between ecological accuracy and feasibility. Some level of coarseness and simplification should be accepted if responses are ecologically justified, and if they can be revised as the knowledge improves. Ideally, response estimates would be provided and updated in a centralized manner by, for example, national environmental administrators, so that different actors would not need to invest on acquiring response estimates case by case.

The responses require ecological attributes to assess the target habitat types’ condition, which were not earlier available for all habitat types in Finland. Here, condition assessment matrices (Supplementary 3) focus on the vegetation and other local ecological attributes specific to each habitat type. Similar condition-based metrics are commonly used in other regions and studies (Borges-Matos et al., 2023; Marshall et al., 2024). In Finland, initial condition is assessed separately for each ecological attribute in each habitat type, and thus the condition of different sites is only comparable within a habitat type. Finnish offset policy specifies which habitat types can be offset with each other (Decree on Voluntary Ecological Offsetting; Suvantola et al. 2024). Most offsets are expected to focus on habitat types in Finland, while offsets for single species may be an exception (e.g., strictly protected species; Suvantola et al. 2024). In such cases, target species require unique ecological attributes and response, as ecological attributes defined for habitat types can poorly represent condition for species (Marshall et al., 2021; Simpson et al., 2022). Utilization of remote sensing products for Finnish condition assessments should also be developed (Mokany et al., 2025). We provided only two options for individual ecological attributes’ weights (1 and 2) and providing a less simplistic weighting scheme might later be appropriate. Overall, the current condition matrices and response estimates work as a baseline that should be tested and revised when needed.

Here we demonstrate the use of the REHAB framework for offsetting, but it can be utilized in other contexts as well. In the EU, the recent Nature Restoration Regulation requires Member States to carry out effective restoration in all directive habitat types (European Parliament 2024). Effectiveness cannot be evaluated without understanding the expected gains and uncertainties of the restoration action; that is, a response estimate. The responses provided here (Supplementary 4) focus on environments and habitats found in Finland, but the approach is applicable to other countries and regions as well. Many of our responses could be applied in the boreal zone of the whole northern Europe. In addition to national restoration commitments, there is a growing interest towards private nature credit markets (Swinfield et al., 2024; Wauchope et al., 2024). Nature credits should be credible, measurable, and verifiable, so that their accumulation can be monitored (IAPB 2024; Wauchope et al., 2024; European Commission 2025). We argue that the REHAB framework as operationalised here could be used as a basis of defining the biodiversity gains that can be used as credible nature credits. Indeed, it would be ideal that in biodiversity offsetting and in the nature credit markets, the units would be the same (habitat hectares) as this would allow the credit markets to be used for purchasing the needed offsets.

Our work provides an important example of successful science–policy-implementation. The presented ecological attributes and their condition matrices have been adopted in the Finnish legislation as the mandatory assessment criteria (Decree on Voluntary Ecological Offsetting, Annex I), without any initial official assignment. Furthermore, the condition assessment matrices have already been used in several environmental and land-use impact assessments. Some of the key factors contributing to the successful implementation were timing, deliberation with a wide range of experts, and an iterative process design. The formulation of condition matrices, offset actions and respective responses occurred at the same time as the national offsetting legislation was finalized and in close communication with the Finnish Ministry of the Environment, which resulted in many concepts from the REHAB approach getting included in the law. During the process, the extensive and diverse expert pool across institutions created a mutual understanding about the REHAB approach, strengthening the outcome and its legitimacy. As a practical result, we were able to gather existing knowledge and practices and avoid unnecessarily creating completely new ones. For example, we used the existing national and EU habitat type evaluation criteria, familiar to at least some experts, as the baseline material for the ecological attributes and their condition matrices, and many attributes ended up utilizing the current practices almost as such. Also, the iterative process between the experts and researchers followed the principles of knowledge co-production (Norström et al., 2020), improving the credibility of the outcome. Especially the in-person workshops were considered valuable for formulating a shared goal and examining key issues via open dialogue, whereas the online workshops and email iteration rounds helped with joint sensemaking.

The strength of offsetting is the clearly defined aim of NNL in a quantifiable manner (Maron et al., 2025). The REHAB approach is a means to achieve this aim in an ecologically realistic way. If one desires ecological accuracy, offsetting becomes inherently complex (Moilanen & Kotiaho, 2018). This fact further highlights the importance of avoiding biodiversity losses by halting conversion of natural land and reducing the consumption of natural resources – a common pledge from offsetting researchers. After all, biodiversity crisis is driven by poor accounting of non-human nature in human decision-making. To overcome this crisis, we humans need to rehabilitate ourselves from the idea that we can proceed without considering the ecological consequences of our actions, whether negative or positive.

## Acknowledgements

We thank the 111 experts who defined the condition metrics and responses. This work was funded by the Strategic Research Council associated with the Academy of Finland (#364448 #364450, #364453, “BOOST”) and the Finnish Ministry of Environment.

## Notes

### Competing Interest Statement

The authors have declared no competing interest.

## References

Airaksinen O, Karttunen K. 2001, Natura 2000-luontotyyppiopas. Ympäristöopas 46. The Finnish environment institute, Helsinki.

BBOP (2012) Standard on Biodiversity Offsets. Washington, D.C., USA: Business and Biodiversity Offsets Programme. https://www.forest-trends.org/wp-content/uploads/imported/BBOP_Standard_on_Biodiversity_Offsets_1_Feb_2013.pdf

Borges-Matos, C., Maron, M., & Metzger, J. P. (2023). A Review of Condition Metrics Used in Biodiversity Offsetting. Environmental Management, 72(4), 727–740. 10.1007/s00267-023-01858-1

Bull, J. W., Lloyd, S. P., & Strange, N. (2017). Implementation Gap between the Theory and Practice of Biodiversity Offset Multipliers. In Conservation Letters (Vol. 10, Number 6, pp. 656–669). Wiley-Blackwell. 10.1111/conl.12335

Bull, J. W., Milner-Gulland, E. J., Addison, P. F. E., Arlidge, W. N. S., Baker, J., Brooks, T. M., Burgass, M. J., Hinsley, A., Maron, M., Robinson, J. G., Sekhran, N., Sinclair, S. P., Stuart, S. N., zu Ermgassen, S. O. S. E., & Watson, J. E. M. (2020). Net positive outcomes for nature. In Nature Ecology and Evolution (Vol. 4, Number 1, pp. 4–7). Nature Research. 10.1038/s41559-019-1022-z

Bull, J. W., Suttle, K. B., Gordon, A., Singh, N. J., & Milner-Gulland, E. J. (2013). Biodiversity offsets in theory and practice. In ORYX (Vol. 47, Number 3, pp. 369–380). 10.1017/S003060531200172X

Chazdon, R. L., Falk, D. A., Banin, L. F., Wagner, M., J. Wilson, S., Grabowski, R. C., & Suding, K. N. (2024). The intervention continuum in restoration ecology: rethinking the active–passive dichotomy. In Restoration Ecology (Vol. 32, Number 8). John Wiley and Sons Inc. 10.1111/rec.13535

Contos, P., Gorrod, E., Caves, K., Oliver, I., & Dorrough, J. W. (2025). How variation among field assessments can affect biodiversity offset outcomes. Conservation Science and Practice. 10.1111/csp2.70096

Curran, M., Hellweg, S., & Beck, J. (2014). Is there any empirical support for biodiversity offset policy? Ecological Applications, 24(4), 617–632. 10.1890/13-0243.1

Decree on Voluntary Ecological Offsetting 933/2023. Finnish Ministry of the Environment. Helsinki.

Defra. 2024. Statutory biodiversity metric tools and guides. GOV.UK. https://www.gov.uk/government/publications/statutory-biodiversity-metric-tools-and-guides

Elo, M., Kareksela, S., Haapalehto, T., Vuori, H., Aapala, K., & Kotiaho, J. S. (2016). The mechanistic basis of changes in community assembly in relation to anthropogenic disturbance and productivity. Ecosphere, 7(4). 10.1002/ecs2.1310

Elo, M., Kareksela, S., Ovaskainen, O., Abrego, N., Niku, J., Taskinen, S., Aapala, K., & Kotiaho, J. S. (2024). Restoration of forestry-drained boreal peatland ecosystems can effectively stop and reverse ecosystem degradation. Communications Earth and Environment, 5(1). 10.1038/s43247-024-01844-3

European Parliament. Nature restoration—European Parliament legislative resolution of 27 February 2024 on the proposal for a regulation of the European Parliament and of the Council on nature restoration (COM(2022)0304—C9-0208/2022-2022/0195(COD)) (European Union, 2024).

Haapalehto, T., Kotiaho, J. S., Matilainen, R., & Tahvanainen, T. (2014). The effects of long-term drainage and subsequent restoration on water table level and pore water chemistry in boreal peatlands. Journal of Hydrology, 519(PB), 1493–1505. 10.1016/j.jhydrol.2014.09.013

Halme, P., Allen, K. A., Auniņš, A., Bradshaw, R. H. W., Brumelis, G., Čada, V., Clear, J. L., Eriksson, A. M., Hannon, G., Hyvärinen, E., Ikauniece, S., Iršenaite, R., Jonsson, B. G., Junninen, K., Kareksela, S., Komonen, A., Kotiaho, J. S., Kouki, J., Kuuluvainen, T., … Zin, E. (2013). Challenges of ecological restoration: Lessons from forests in northern Europe. In Biological Conservation (Vol. 167, pp. 248–256). 10.1016/j.biocon.2013.08.029

Hemming, V., Burgman, M. A., Hanea, A. M., McBride, M. F., & Wintle, B. C. (2018). A practical guide to structured expert elicitation using the IDEA protocol. Methods in Ecology and Evolution, 9(1), 169–180. 10.1111/2041-210X.12857

IAPBB (2024) Framework for high integrity biodiversity credit markets. International Advisory Panel on Biodiversity Credits. https://www.iapbiocredits.org/framework

IUCN (2016) IUCN Policy on Biodiversity Offsets. International Union for Conservation of Nature. https://portals.iucn.org/library/sites/library/files/resrecfiles/WCC_2016_RES_059_EN.pdf

Kangas, J., Kullberg, P., Pekkonen, M., Kotiaho, J. S., & Ollikainen, M. (2021). Precision, Applicability, and Economic Implications: A Comparison of Alternative Biodiversity Offset Indexes. Environmental Management, 68(2), 170–183. 10.1007/s00267-021-01488-5

Kontula T, Raunio A. 2019. Threatened habitat types in Finland 2018 - Red List of habitats results and basis for assessment. The Finnish Environment 5/2018.

Laitila, J., Moilanen, A., & Pouzols, F. M. (2014). A method for calculating minimum biodiversity offset multipliers accounting for time discounting, additionality and permanence. Methods in Ecology and Evolution, 5(11), 1247–1254. 10.1111/2041-210X.12287

Maron, M., Hobbs, R. J., Moilanen, A., Matthews, J. W., Christie, K., Gardner, T. A., Keith, D. A., Lindenmayer, D. B., & McAlpine, C. A. (2012). Faustian bargains? Restoration realities in the context of biodiversity offset policies. Biological Conservation, 155, 141–148. 10.1016/j.biocon.2012.06.003

Maron, M., Ives, C. D., Kujala, H., Bull, J. W., Maseyk, F. J. F., Bekessy, S., Gordon, A., Watson, J. E. M., Lentini, P. E., Gibbons, P., Possingham, H. P., Hobbs, R. J., Keith, D. A., Wintle, B. A., & Evans, M. C. (2016). Taming a Wicked Problem: Resolving Controversies in Biodiversity Offsetting. In BioScience (Vol. 66, Number 6, pp. 489–498). Oxford University Press. 10.1093/biosci/biw038

Maron, M., von Hase, A., Quétier, F., Sonter, L. J., Theis, S., & zu Ermgassen, S. O. S. E. (2025). Biodiversity offsets, their effectiveness and their role in a nature positive future. Nature Reviews Biodiversity, 1(3), 183–196. 10.1038/s44358-025-00023-2

Marshall, E., Southwell, D., Wintle, B. A., & Kujala, H. (2024). A global analysis reveals a collective gap in the transparency of offset policies and how biodiversity is measured. In Conservation Letters (Vol. 17, Number 1). John Wiley and Sons Inc. 10.1111/conl.12987

Marshall, E., Valavi, R., Connor, L. O., Cadenhead, N., Southwell, D., Wintle, B. A., & Kujala, H. (2021). Quantifying the impact of vegetation-based metrics on species persistence when choosing offsets for habitat destruction. Conservation Biology, 35(2), 567–577. 10.1111/cobi.13600

Martin, T. G., Burgman, M. A., Fidler, F., Kuhnert, P. M., Low-Choy, S., Mcbride, M., & Mengersen, K. (2012). Eliciting Expert Knowledge in Conservation Science. In Conservation Biology (Vol. 26, Number 1, pp. 29–38). 10.1111/j.1523-1739.2011.01806.x

Maseyk, F. J. F., Barea, L. P., Stephens, R. T. T., Possingham, H. P., Dutson, G., & Maron, M. (2016). A disaggregated biodiversity offset accounting model to improve estimation of ecological equivalency and no net loss. Biological Conservation, 204, 322–332. 10.1016/j.biocon.2016.10.016

Moilanen, A., Jalkanen, J., Halme, P., Nieminen, E., Kotiaho, J. S., & Kujala, H. (2024). Monitoring in biodiversity offsetting. In Global Ecology and Conservation (Vol. 54, p. e03039). Elsevier B.V. 10.1016/j.gecco.2024.e03039

Moilanen, A., & Kotiaho, J. S. (2018). Fifteen operationally important decisions in the planning of biodiversity offsets. In Biological Conservation (Vol. 227, pp. 112–120). Elsevier Ltd. 10.1016/j.biocon.2018.09.002

Moilanen, A., & Lehtinen, P. (2025). Simple analysis of biodiversity response functions and multipliers for biodiversity offsetting and other applications. Environmental Modelling and Software, 185, 106322. 10.1016/j.envsoft.2025.106322

Mokany, K., Ware, C., Valavi, R., Giljohann, K., Ferrier, S., Stitzlein, C., & Mata, G. (2025). A habitat-based approach to reporting the direct impacts of an organization on biodiversity. Conservation Biology, e70071. 10.1111/cobi.70071

Norström, A. V., Cvitanovic, C., Löf, M. F., West, S., Wyborn, C., Balvanera, P., Bednarek, A. T., Bennett, E. M., Biggs, R., de Bremond, A., Campbell, B. M., Canadell, J. G., Carpenter, S. R., Folke, C., Fulton, E. A., Gaffney, O., Gelcich, S., Jouffray, J. B., Leach, M., … Österblom, H. (2020). Principles for knowledge co-production in sustainability research. Nature Sustainability, 3(3), 182–190. 10.1038/s41893-019-0448-2

Prober, S. M., Liedloff, A. C., England, J. R., Mokany, K., Ogilvy, S., & Richards, A. E. (2025). Accounting for the biodiversity benefits of woody plantings in agricultural landscapes: A global meta-analysis. In Agriculture, Ecosystems and Environment (Vol. 381). Elsevier B.V. 10.1016/j.agee.2024.109453

Ramberg, E., Berglund, H., Penttilä, R., Strengbom, J., & Jönsson, M. (2023). Prescribed fire is an effective restoration measure for increasing boreal fungal diversity. Ecological Applications, 33(6). 10.1002/eap.2892

Shorohova, E., Lindberg, H., Kuuluvainen, T., & Vanha-Majamaa, I. (2024). Deadwood enrichment in Fennoscandian spruce forests – New results from the EVO experiment. Forest Ecology and Management, 564. 10.1016/j.foreco.2024.122013

Siitonen, J., Penttilä, R., & Ihalainen, A. (2012). METSO-ohjelman uusien pysyvien ja määräaikaisten suojelualueiden ekologinen laatu Uudenmaan alueella. Metsätieteen Aikakausikirja, 4, 259–284.

Simpson, K. H., de Vries, F. P., Dallimer, M., Armsworth, P. R., & Hanley, N. (2022). Ecological and economic implications of alternative metrics in biodiversity offset markets. Conservation Biology, 36(5). 10.1111/cobi.13906

Soomets, E., Lõhmus, A., & Rannap, R. (2023). Restoring functional forested peatlands by combining ditch-blocking and partial cutting: An amphibian perspective. Ecological Engineering, 192. 10.1016/j.ecoleng.2023.106968

Spake, R., Ezard, T. H. G., Martin, P. A., Newton, A. C., & Doncaster, C. P. (2015). A meta-analysis of functional group responses to forest recovery outside of the tropics. Conservation Biology, 29(6), 1695–1703. 10.1111/cobi.12548

Suvantola, L., Borgström, S., Härkönen, S. & Ojala, O. (2024) Voluntary Ecological Offsetting – Guidance document. Ministy of Environment 12/2024. https://ym.fi/documents/1410903/0/Guidance+document+for+voluntary+ecological+offsetting+in+FInland.pdf/1175fa1b-7f5f-7b8d-a1d5-680cf7e4e843/Guidance+document+for+voluntary+ecological+offsetting+in+FInland.pdf?t=1734504565921

Strange, N., Ermgassen, S. zu, Marshall, E., Bull, J. W., & Jacobsen, J. B. (2024). Why it matters how biodiversity is measured in environmental valuation studies compared to conservation science. Biological Conservation, 292. 10.1016/j.biocon.2024.110546

Swinfield, T., Shrikanth, S., Bull, J. W., Madhavapeddy, A., & zu Ermgassen, S. O. S. E. (2024). Nature-based credit markets at a crossroads. In Nature Sustainability. Nature Research. 10.1038/s41893-024-01403-w

Syrjänen, K., Hakalisto, S., Mikkola, J., Musta, I., Nissinen, M., Savolainen, R., Seppälä, J., Seppälä, M., Siitonen, J., & Valkeapää, A. (2016). Monimuotoisuudelle arvokkaiden metsäelinympäristöjen tunnistaminen: METSO ohjelman luonnontieteelliset valintaperusteet 2016–2025. Ympäristöministeriön Raportteja 17/2016.

Toivonen, M., Herzon, I., & Kuussaari, M. (2015). Differing effects of fallow type and landscape structure on the occurrence of plants, pollinators and birds on environmental fallows in Finland. Biological Conservation, 181, 36–43. 10.1016/j.biocon.2014.10.034

Wauchope, H. S., zu Ermgassen, S. O. S. E., Jones, J. P. G., Carter, H., to Bühne, H. S., & Milner-Gulland, E. J. (2024). What is a unit of nature? Measurement challenges in the emerging biodiversity credit market. In Proceedings of the Royal Society B: Biological Sciences (Vol. 291, Number 2036). Royal Society Publishing. 10.1098/rspb.2024.2353

zu Ermgassen, S. O. S. E., Baker, J., Griffiths, R. A., Strange, N., Struebig, M. J., & Bull, J. W. (2019). The ecological outcomes of biodiversity offsets under “no net loss” policies: A global review. In Conservation Letters (Vol. 12, Number 6). Wiley-Blackwell. 10.1111/conl.12664

zu Ermgassen, S. O. S. E., Marsh, S., Ryland, K., Church, E., Marsh, R., & Bull, J. W. (2021). Exploring the ecological outcomes of mandatory biodiversity net gain using evidence from early-adopter jurisdictions in England. Conservation Letters, 14(6). 10.1111/conl.12820

