## Supplement 1-2 for "Defining ecologically realistic biodiversity offset multipliers with the Response-based Habitat Hectare Assessment of Biodiversity Gains (REHAB)"

**Supplementary 1 – Calculating response functions**

Restoration and passive recovery

The parameters required for active restoration measures and passive recovery:

- *c_0_*: Minimum initial condition. The lowest condition, where the action is still appropriate for the habitat type.
- *c_max_*: Final condition. The maximum condition that the target habitat type can eventually reach when the action is conducted to in the minimum initial condition (*c_max_*).
- *T*: Duration. The number of years the target habitat type takes to develop from the minimum initial (*c_0_*) to the final condition (*c_max_*).
- *u*: Uncertainty value (%).

First, the proportion of the duration, *p(t)*, is calculated, that describes the proportion of the year *t* from the total response duration *T*. After *T* years, the habitat type has reached its final condition, and the proportion of the duration will be set 1:

$$p\left( t \right)=\left\{ \begin{aligned} 1, t\geq T \\ \frac{t}{T}, t<T \end{aligned} \right.$$

Then, the condition improvement, i.e., the response value, at year *t*, *i(t)*, is calculated as follows depending on the shape of the response function (Figure S1.1):

- Convex, less steep: $i\left( t \right)={p(t)}^{0.5}(c_{max}-c_{0})$
- Convex, steeper: $i\left( t \right)={p(t)}^{0.33}(c_{max}-c_{0})$
- Concave, less steep: $i\left( t \right)=\left( \frac{1}{2^{3}} \right)\left( c_{max}-c_{0} \right)\left( \frac{p\left( t \right)}{0.5} \right)^{3}$
- Concave, steeper: $i\left( t \right)=\left( \frac{1}{2^{1.5}} \right)\left( c_{max}-c_{0} \right)\left( \frac{p\left( t \right)}{0.5} \right)^{1.5}$
- Linear: $i\left( t \right)=p\left( t \right)(c_{max}-c_{0})$
- Sigmoid: $i\left( t \right)=\frac{\left( c_{max}-c_{0} \right)}{2}+\left( c_{max}-c_{0} \right)\frac{\left( p\left( t \right)-0.5 \right)^{\frac{1}{3}}}{2\times{0.5}^{\frac{1}{3}}}$

Finally, uncertainty-corrected response, *i_u_(t)* is calculated by multiplying the response with the certain proportion of the response estimate (i.e., 1 – uncertainty):

$$i\left( t \right)=i(t)\times(1-\frac{u}{100})$$

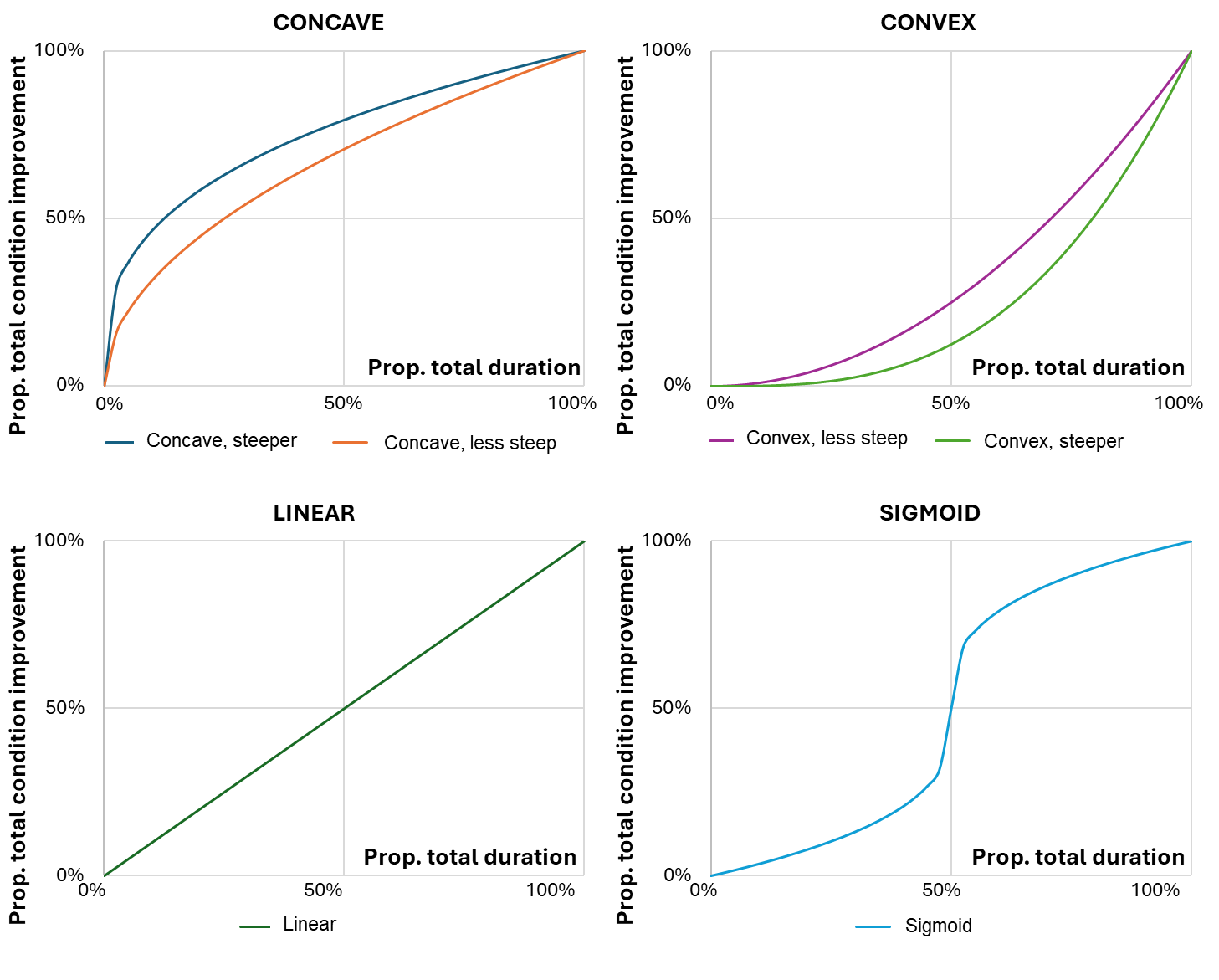

**Figure S1.1.** Different response function options for restoration and passive recovery. The x-axis shows the proportion of the response-specific duration, and the y-axis shows the proportion of the response-specific total improvement (final condition minus lowest condition).

Management

Calculating biodiversity gains from management requires condition responses with and without management. Here, only baseline response without management is calculated. For simplicity, a linear decrease in condition is assumed, until the habitat type reaches its final condition. Furthermore, for simplicity, the highest possible condition 1 hha/ha can be assumed as the starting point.

The required parameters are:

- *c_min_*: Final condition. The condition the habitat type eventually ends up in when left without management.
- *T*: Duration. The number of years it takes to reach the final condition, assuming initial condition of 1 hha/ha.

The proportion of the duration, *p(t)*, is calculated as above. The condition at the year *t*, *c(t)*, is then calculated as follows (Figure S1.2.):

$$c\left( t \right)=1-(p\left( t \right)\left( 1-c_{min} \right))$$

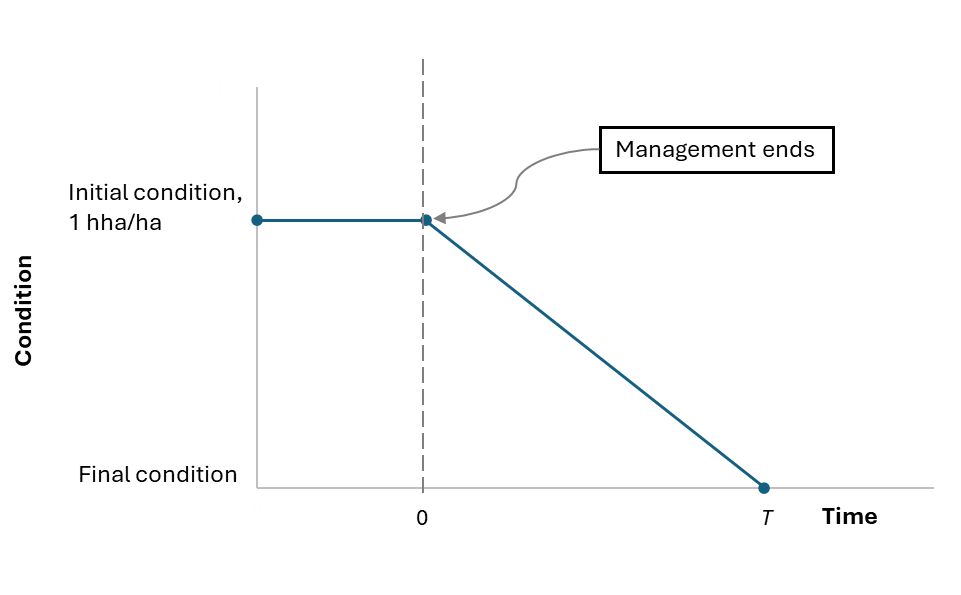

**Figure S1.2.** Condition response without management.

Furthermore, the yearly loss in condition, *l*, can be calculated that allows predicting future condition without management, and how rapidly the lowest possible condition is reached. If the initial condition of a site is below habitat type-specific lowest final condition, mere management is not considered to suffice, but active restoration measures are also needed.

The yearly loss in condition, *l*, is calculated as:

$$l=\frac{\left( 1-c_{min} \right)}{T}$$

**Supplementary 2 – Geographical regions**

The experts were asked to define the duration of responses separately for southern and northern Finland (Fig. S2.1.) for the habitats that span across the regions. The marine (Baltic Sea) and coastal habitats only exist in the southern, and fell habitats in the northern region.

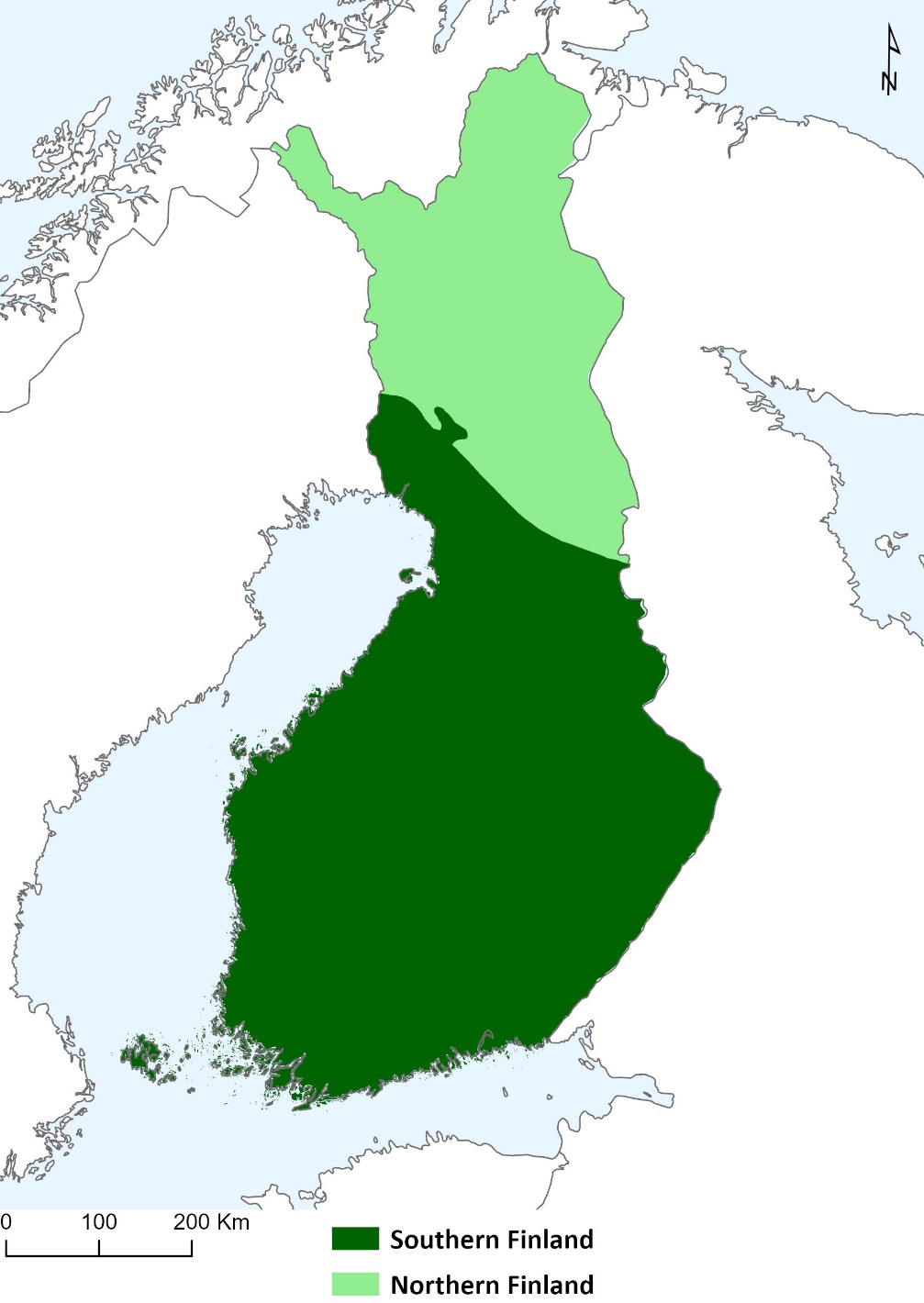

**Figure S2.1.** Southern and northern Finland.
