## Supplement 3 for "Defining ecologically realistic biodiversity offset multipliers with the Response-based Habitat Hectare Assessment of Biodiversity Gains (REHAB)"

Condition assessment matrices for Finnish habitat types (versions 1–2025).

These assessment matrices are meant to guide condition assessment of habitat types in the field. These materials are written for professionals. Each ecological attribute is compared against pristine habitat types. Attributes are estimated onsite; no exact measurement nor species inventories are required. One can choose a condition class for each attribute that has no verbal description; the descriptions are meant to help field assessor to “calibrate their eyes” in the field. Detailed assessment guides for each ecological attribute for each assessment matrix are provided in Finnish.

The condition of a habitat patch is the weighted mean of the habitat type specific attribute scores. Thus, a score for each attribute is required. When uncertain about which condition class to choose for a given ecological attribute, one should select the higher option.

### 1. The Baltic Sea

#### 1.1. Hard benthic habitats characterized by perennial algae or aquatic moss, Hard benthic habitats characterized by invertebrates, Benthic habitats characterized by filamentous annual algae, Benthic habitats characterized by microphytobenthic organisms and grazing snails, and other habitat types found on hard substrates

|  | **(Primary)**  **Benthos representativeness: community and community structure (abundance, density, coverage, biomass, species/species complex abundance ratios)** | **(Primary)**  **Epiphytic algae, quantity of loose sediment^1^** | **(Secondary)**  **Secchi depth^2^** | **(Secondary)**  **Sessile benthic invasive alien species** | **(Secondary)**  **Aquatic construction and benthic modification (e.g. dredging, piling), other anthropogenic pressures** |
| --- | --- | --- | --- | --- | --- |
| **Relative attribute weight** | **2** | **2** | **1** | **1** | **1** |
| **1.0 (EXCELLENT)** | Representative benthos.  Community and community structure are characteristic of habitat type, geographical area, and local conditions. No signs of altered community or community structure e.g. due to eutrophication. | No epiphytic algae or sediments. | Excellent. | No invasive alien species. | Benthic shows no signs of aquatic construction or modification. |
| **0.9** |  |  |  |  |  |
| **0.8** |  |  |  |  |  |
| **0.7 (GOOD)** |  | Low levels of epiphytic algae and/or sediments. | Good. |  | Benthic shows signs of construction and/or modification (e.g. dredging), but community characteristic of habitat type has recovered. |
| **0.6** |  |  |  |  |  |
| **0.5 (SATISFACTORY)** | Benthos representativeness decreased. Some species characteristic of habitat type are present, but community and/or community structure are altered. | Moderate levels of epiphytic algae and/or sediments. | Satisfactory. | Invasive alien species are present, but current levels do not impede the structure or functioning of habitat type. However, a distinct risk exists that levels will increase to impeding. |  |
| **0.4** |  |  |  |  |  |
| **0.3 (POOR)** |  |  | Tolerable. |  | Benthic has been constructed or modified, but community characteristic of habitat type shows signs of recovery. |
| **0.2** |  |  |  |  |  |
| **0.1 (VERY POOR)** | Unrepresentative benthos. Community highly altered, and/or species characteristic of habitat type and site are barely present. Community structure not characteristic of habitat type. | Abundant epiphytic algae and/or sediment. | Poor. | Structure and/or functioning of habitat type is clearly reduced due to invasive alien species. | Benthic is greatly modified (e.g. dredged, blasted) or is completely constructed, coated, or artificial and does not uphold community characteristic of habitat type, community cannot recover in current conditions. Pronounced erosion or extensive water traffic may also be present (e.g. a harbour). |
| **0.0 (Not a habitat type)** |  | | | | |

^1^Sediment quantity: see VELMU survey instructions, Finnish Environment Institute & Metsähallitus (Finland) 2022. The Finnish Inventory Programme for the Underwater Marine Environment (VELMU) Methodology guide. Version 14.02.2022. In Finnish.

^2^Secchi depth range: Status classification and assessment criteria of surface waters in the third river basin management cycle. Finnish Environment Institute SYKE reports 37/2019, Annex 9. In Finnish with English summary.

#### 1.2. Reefs and sandbanks

|  | **(Primary)**  **Benthos representativeness: community and community structure (abundance, density, coverage, biomass, species/species complex abundance ratios)** | **(Primary)**  **Epiphytic algae, quantity of loose sediment^1^** | **(Secondary)**  **Secchi depth^2^** | **(Secondary)**  **Aquatic construction and benthic modification (e.g. dredging, piling), other anthropogenic pressures** |
| --- | --- | --- | --- | --- |
| **Relative attribute weight** | **2** | **1** | **1** | **1** |
| **1.0 (EXCELLENT)** | Representative benthos.  Community and community structure are characteristic of habitat type, geographical area, and local conditions. No signs of altered community or community structure e.g. due to eutrophication. | No epiphytic algae or sediments. | Excellent. | Benthic shows no signs of aquatic construction or modification. |
| **0.9** |  |  |  |  |
| **0.8** |  |  |  |  |
| **0.7 (GOOD)** |  | Low levels of epiphytic algae and/or sediments. | Good. | Benthic shows signs of construction and/or modification (e.g. dredging), but community characteristic of habitat type has recovered. |
| **0.6** |  |  |  |  |
| **0.5 (SATISFACTORY)** | Benthos representativeness decreased. Some species characteristic of habitat type are present, but community and/or community structure are altered. | Moderate levels of epiphytic algae and/or sediments. | Satisfactory. |  |
| **0.4** |  |  |  |  |
| **0.3 (POOR)** |  |  | Tolerable. | Benthic has been constructed or modified, but community characteristic of habitat type shows signs of recovery. |
| **0.2** |  |  |  |  |
| **0.1 (VERY POOR)** | Unrepresentative benthos. Community is highly altered, and/or species characteristic of habitat type and site are barely present. Community structure is not characteristic of habitat type. | Abundant epiphytic algae and/or sediment. | Poor. | Benthic is greatly modified (e.g. dredged, blasted) or is completely constructed, coated, or artificial and does not uphold community characteristic of habitat type, community cannot recover in current conditions. Pronounced erosion or extensive water traffic may also occur (e.g. a harbour). |
| **0.0 (Not a habitat type)** |  | | | |

^1^Sediment quantity: see VELMU survey instructions, Finnish Environment Institute & Metsähallitus (Finland) 2022. The Finnish Inventory Programme for the Underwater Marine Environment (VELMU) Methodology guide. Version 14.02.2022. In Finnish.

^2^Secchi depth reference values: Status classification and assessment criteria of surface waters in the third river basin management cycle. Finnish Environment Institute SYKE reports 37/2019, annex 9. In Finnish with English summary.

#### 1.3. Soft benthic habitats characterized by vegetation

|  | **(Primary)**  **Benthos representativeness: community and community structure (abundance, density, coverage, biomass, species/species complex abundance ratios)** | **(Primary)**  **Epiphytic algae, quantity of loose sediment^1^** | **(Secondary)**  **Secchi depth^2^** | **(Secondary)**  **Overgrowth caused by helophytes and floating-leaved plants that is not natural succession** | **(Secondary)**  **Sessile benthic invasive alien species** | **(Secondary)**  **Aquatic construction and benthic modification (e.g. dredging, piling), other anthropogenic pressures** | **(Secondary)**  **Additionally, in the benthic of shallow sea bays: impact of surrounding watershed** |
| --- | --- | --- | --- | --- | --- | --- | --- |
| **Relative attribute weight** | **2** | **1** | **1** | **1** | **1** | **1** | **1** |
| **1.0 (EXCELLENT)** | Representative benthos.  Community and community structure are characteristic of habitat type, geographical area, and local conditions. No signs of altered community or community structure e.g. due to eutrophication. | No epiphytic algae or sediments. | Excellent. | No overgrowth caused by helophytes or floating-leaved plants is present or overgrowth is not accelerating. | No invasive alien species present. | Benthic shows no signs of aquatic construction or modification. | Watershed is pristine or near-natural, and loading from watershed does not surpass natural washout (e.g. nutrients, solid particles, humus). |
| **0.9** |  |  |  |  |  |  |  |
| **0.8** |  |  |  |  |  |  |  |
| **0.7 (GOOD)** |  | Low levels of epiphytic algae and/or sediments present. | Good. | Overgrowth caused by helophytes or floating-leaved plants is present, but overgrowth has not been particularly rapid (verifiable e.g. using aerial photographs), or reed encroachment likely does threaten the entire benthic habitat type. |  | Benthic shows signs of construction and/or modification (e.g. dredging), but community characteristic of habitat type has recovered. |  |
| **0.6** |  |  |  |  |  |  |  |
| **0.5 (SATISFACTORY)** | Benthos representativeness decreased. Some species characteristic of habitat type are present, but community and/or community structure are altered. | Moderate levels of epiphytic algae and/or sediments. | Satisfactory. |  | Invasive alien species are present, but currently not at levels that impede the structure or functioning of habitat type. However, a distinct risk exists that levels will increase to impeding. |  | Watershed has been modified, and/or loading from watershed certainly or likely surpasses natural washout. |
| **0.4** |  |  |  |  |  |  |  |
| **0.3 (POOR)** |  |  | Tolerable. | Clear overgrowth caused by helophytes- or floating-leaved plants, overgrowth is rapid (verifiable e.g. using aerial photographs), or complete benthic layer overgrowth is possible (e.g. due to shallow conditions or peat formation. |  | Benthic has been constructed or modified, but community characteristic of habitat type shows signs of recovery. |  |
| **0.2** |  |  |  |  |  |  |  |
| **0.1 (VERY POOR)** | Unrepresentative benthos. Community is highly altered, and/or species characteristic of habitat type and site are barely present. Community structure uncharacteristic of habitat type. | Abundant epiphytic algae and/or sediment. | Poor. | Fully overgrown. | Structure and/or functioning of habitat type are clearly reduced due to invasive alien species. | Benthic is greatly modified (e.g. dredged, blasted) or is completely constructed, coated, or artificial and does not uphold community characteristic of habitat type, community cannot recover in current conditions. Pronounced erosion or extensive water traffic may als occur (e.g. a harbour). | Watershed is mostly constructed/modified (e.g. population centres, agriculture), and/or loading from the watershed is certainly or likely very pronounced, transgressing natural washout. |
| **0.0 (Not a habitat type)** |  | | | | | | |

^1^Sediment quantity: see VELMU survey instructions, Finnish Environment Institute & Metsähallitus (Finland) 2022. The Finnish Inventory Programme for the Underwater Marine Environment (VELMU) Methodology guide. Version 14.02.2022. In Finnish.

^2^Secchi depth reference values: Status classification and assessment criteria of surface waters in the third river basin management cycle. Finnish Environment Institute SYKE reports 37/2019, annex 9. In Finnish with English summary.

#### 1.4. Hard benthic habitats characterized by invertebrates, and other habitat types found on soft substrates

|  | **(Primary)**  **Benthos representativeness: community and community structure (abundance, density, coverage, biomass, species/species complex abundance ratios)** | **(Primary)**  **Benthic oxygen deficit** | **(Secondary)**  **Aquatic construction and benthic modification (e.g. dredging, piling), other anthropogenic pressures** |
| --- | --- | --- | --- |
| **Relative attribute weight** | **2** | **2** | **1** |
| **1.0 (EXCELLENT)** | Representative benthos.  Community and community structure are characteristic of habitat type, geographical area, and local conditions. No signs of altered community or community structure e.g. due to eutrophication. | Oxygen concentration surpasses 8 mg/l. | Benthic shows no signs of aquatic construction or modification. |
| **0.9** |  |  |  |
| **0.8** |  |  |  |
| **0.7 (GOOD)** |  | Oxygen concentration 6–8 mg/l. | Benthic shows signs of construction and/or modification (e.g. dredging), but community characteristic of habitat type has recovered. |
| **0.6** |  |  |  |
| **0.5 (SATISFACTORY)** | Benthos representativeness decreased. Some species characteristic of habitat type are present, but community and/or community structure are altered. | Oxygen concentration 4–6 mg/l. |  |
| **0.4** |  |  |  |
| **0.3 (POOR)** |  | Oxygen concentration 2–4 mg/l. | Benthic has been constructed or modified, but community characteristic of habitat type shows signs of recovery. |
| **0.2** |  |  |  |
| **0.1 (VERY POOR)** | Unrepresentative benthos. Community is highly altered, and/or species characteristic of habitat type and site are barely present. Community structure is not characteristic of habitat type. | Oxygen concentration 0–2 mg/l. | Benthic is greatly modified (e.g. dredged, blasted) or is completely constructed, coated, or artificial and does not uphold community characteristic of habitat type, community cannot recover in current conditions. Pronounced erosion or extensive water traffic may also be present (e.g. a harbour). |
| **0.0 (Not a habitat type)** |  | | |

#### 1.5. Flada-lakes (coastal lagoons), Glo-lakes (coastal lagoons), Coastal estuaries

|  | **(Primary)**  **Vegetation representativeness: plant community and community structure (abundance, coverage, species/species complex abundance ratios)** | **(Secondary)**  **Impact of surrounding watershed** | **(Secondary)**  **Modification, construction, and other anthropogenic pressure** | **(Primary)**  **Additionally for flada-lakes: sill** |
| --- | --- | --- | --- | --- |
| **Relative attribute weight** | **2** | **1** | **1** | **Flada-lakes 2, not applicable to other types.** |
| **1.0 (EXCELLENT)** | Representative vegetation. Community and vegetation structure are characteristic and representative of habitat type, geographical area, local conditions, and natural successional stage. Community exhibits small-scale variation and transitioning between various benthic habitat types and between underwater and shoreline habitats. No alien species present. | Watershed is pristine or near-natural, and loading from watershed does not surpass natural washout (e.g. nutrients, solid particles, humus). | Site is completely unconstructed and unmodified (e.g. not dredged), and its succession can continue naturally. Virtually no signs of anthropogenic activity, at most individual docks, boats etc. can be present. | Pristine inflow and sill. |
| **0.9** |  |  |  |  |
| **0.8** |  |  |  |  |
| **0.7 (GOOD)** | Vegetation representativeness has decreased to a degree. Essential community characteristic of habitat type is observable but is not as representative as in the ‘excellent’ category. Vegetation structure may be altered to a degree. Small-scale variation occurs in community, but is not as representative as in the ‘Excellent’ category. For example, may exhibit slight signs of eutrophication-induced overgrowth. No alien species present. |  | Site shows signs of modification (e.g. dredging), but community characteristic of habitat type has recovered or is recovering. Minor anthropogenic activity is present, e.g. docks, small boat ramp, or other small-scale construction | Inflow partially dredged, but sill still exists. |
| **0.6** |  |  |  |  |
| **0.5 (SATISFACTORY)** |  | Watershed has been modified, and/or loading from the watershed certainly or likely surpasses natural washout. | Site is mostly unconstructed, but anthropogenic activity is fairly abundant, continuous, or regular (e.g. a port). Natural succession may be hindered, e.g. due to regular dredging. |  |
| **0.4** |  |  |  |  |
| **0.3 (POOR)** | Vegetation representativeness is distinctly decreased. Community and/or vegetation structure exhibit clear changes, some representative species are present. For example, clear eutrophication-induced overgrowth may occur. Alien species may occur. |  |  |  |
| **0.2** |  |  |  |  |
| **0.1 (VERY POOR)** | Vegetation is unrepresentative. Community is greatly altered, and/or species characteristic of habitat type and site are barely present at all, vegetation structure is uncharacteristic of habitat type. May be completely overgrown. May be completely overtaken by alien species. No natural variability and transitioning occur between underwater and shoreline habitats. | Watershed is mostly constructed/modified (e.g. population centres, agriculture), and/or loading from the watershed is certainly or likely very pronounced, transgressing natural washout. | Watershed constructed, modified, or artificial throughout. Intensive anthropogenic activity. Natural succession may be hindered, e.g. due to regular dredging. | Inflow is widely dredged and sill is completely destroyed, or inflow is terraced and connection to sea is completely blocked. |
| **0.0 (Not a habitat type)** |  | | | |

### 2. The Baltic Sea coast

#### 2.1. Coastal habitat types excluding early-successional forests

|  | **(Primary)**  **Vegetation representativeness: (vascular plants and/or lichens and/or mosses): community and vegetation structure (abundance, coverage, species/species complex abundance ratios)** | **(Secondary)**  **Build-up of filamentous algae mass on the shore** | **(Secondary)**  **Invasive alien plants** | **(Secondary)**  **Construction, modification, other anthropogenic impact** | **(Primary)**  **Additionally for coastal wooded dunes: forest structure (uneven-agedness, stochastic spatial distribution, dead wood quantity and continuum, semi-openness, old trees, wind effect)** |
| --- | --- | --- | --- | --- | --- |
| **Relative attribute weight** | **2** | **1** | **1** | **1** | **Coastal wooded dunes 2, not applied to other habitat types** |
| **1.0 (EXCELLENT)** | Representative vegetation. Community and vegetation structure are characteristic and representative of habitat type, geographical area, local conditions, and natural successional stage. | No harmful mass of filamentous algae. | No invasive alien plant species. | No signs or very few signs of anthropogenic activity. Shore completely unconstructed. At most, individual paths or duckboards and light recreational use. | All structural attributes characteristic of habitat type, local conditions, and natural successional stage are present. |
| **0.9** |  |  |  |  |  |
| **0.8** |  |  |  |  |  |
| **0.7 (GOOD)** | Vegetation representativeness is decreased to a degree. Essential community characteristic of habitat type is observable but is not as representative as in the ‘excellent’ category. Vegetation structure may be altered to a degree. |  | Individual invasive alien plants are present. |  | Forest strucure is slightly altered due to anthropogenic activity. |
| **0.6** |  |  |  |  |  |
| **0.5 (SATISFACTORY)** | Vegetation representativeness is distinctly decreased. Community and/or vegetation structure exhibit clear changes, some representative species are present. | Occasional thick filamentous algae coverage or wider coverage of thin filamentous algae on the shoreline. |  | Minor anthropogenic activity. Possible signs of light construction and/or modification that barely impact habitat type (paths, dock, small boat ramp, minor terrain erosion). |  |
| **0.4** |  |  |  |  |  |
| **0.3 (POOR)** |  |  | Several occurrences of invasive alien plants. | Moderate anthropogenic activity. Shore may be constructed or modified in a way that impacts the habitat type and/or its rehabilitation possiblities (e.g. numerous recreational cabins, dredgings, regular traffic, terrain erosion). | Forest structure is clearly altered due to anthropogenic activity. |
| **0.2** |  |  |  |  |  |
| **0.1 (VERY POOR)** | Vegetation is unrepresentative. Community is greatly altered, and/or species characteristic of habitat type and site are barely present, vegetation structure is uncharacteristic of habitat type. | Shore is largely covered in a thick layer of filamentous algae. | Area is largerly taken over by invasive alien plants. | Intensive anthropogenic activity. Shore may be completely constructed, modified, or artificial. Sandy shores may have a swimming beach that is maintained regularly. | Forest structure is uncharacteristic of habitat type. |
| **0.0 (Not a habitat type)** |  | | | | |

#### 2.2. Coastal early-successional forests

|  | **(Primary)**  **Stand structural attributes (multiple species, uneven-agedness, stratification, zonality, stochastic spatial distribution)** | **(Primary)**  **Position in successional series** | **(Secondary)**  **Dead wood**  *Including all dead wood objects* | **(Secondary)**  **Invasive alien** **plants** | **(Secondary)**  **Construction, modification, other anthropogenic impact** |
| --- | --- | --- | --- | --- | --- |
| **Relative attribute weight** | **2** | **2** | **1** | **1** | **1** |
| **1.0 (EXCELLENT)** | All structural attributes characteristic of habitat type, geographical area, local conditions, and natural successional stage are present. | Site is part of a wider pristine or near-natural successional sere and is bordered by a wooded or scrub area in the same series. | Dead wood quantity is characteristic of habitat type, local conditions, and natural successional stage, and natural dead wood generation is undisturbed. | No invasive alien plant species. | No signs or very few signs of anthropogenic activity. Shore is completely unconstructed. At most, individual paths and duckboards, light recreational use, forest grazing. |
| **0.9** |  |  |  |  |  |
| **0.8** |  |  |  |  |  |
| **0.7 (GOOD)** | Forest strucure is slightly altered due to anthropogenic activity. |  | Dead wood is present, but at lower levels than is characteristic for habitat type, local conditions, and natural successional stage, and/or, at most, natural dead wood generation is slightly decreased. | Individual invasive alien plants. |  |
| **0.6** |  |  |  |  |  |
| **0.5 (SATISFACTORY)** |  | Site is part of a wider successional series, but the ecological status of the series is clearly reduced or the series has been interrupted. |  |  | Minor anthropogenic activity. For example, harvester tracks, selection felling, individual forest ditches. Shore can be lightly constructed and modified in a way that barely impacts the habitat type (e.g. dock, small boat ramp). |
| **0.4** |  |  |  |  |  |
| **0.3 (POOR)** | Forest structure is clearly altered due to anthropogenic activity. |  | Dead wood is sparsely spaced out or present as individual objects, and/or natural dead wood generation is clearly decreased. | Several occurrences of invasive alien plants. | Moderate anthropogenic activity. For example, terrain erosion, thinning fellings, and logging residue piles, ditching etc. Shore may be constructed or modified in a way that impacts habitat type (e.g. numerous recreational cabins, dredgings, regular traffic). |
| **0.2** |  |  |  |  |  |
| **0.1 (VERY POOR)** | Forest structure is uncharacteristic of habitat type. | Site is not part of a wider successional series. | No dead wood, and/or natural dead wood generation is greatly decreased. | Area is largerly taken over by invasive alien plants. | Intensive anthropogenic activity. For example, marked terrain erosion, soil tillage, clear-cut fellings. Shore may be completely constructed, modified, or artificial. |
| **0.0 (Not a habitat type)** |  | | | | |

### 3. Inland waters and shores: lakes and ponds

#### 3.1. Lakes to which the ecological status classification of the EU Water Framework Dirctive (WFD) can be applied

|  | **(Primary)**  **WFD: aquatic vegetation** | **(Primary)**  **WFD: Level of hydromorphological change** | **(Primary)**  **WFD: benthos (if applicable and necessary for project impacts)** | **(Primary)**  **Watershed condition** |
| --- | --- | --- | --- | --- |
| **Relative attribute weight** | **1** | **1** | **1** | **1** |
| **1.0 (EXCELLENT)** | Excellent. | Excellent. | Excellent. | Watershed is pristine or near-natural, and loading from watershed does not surpass natural washout (e.g. nutrients, solid particles, humus). |
| **0.9** |  |  |  |  |
| **0.8** |  |  |  |  |
| **0.7 (GOOD)** | Good. | Good. | Good. |  |
| **0.6** |  |  |  |  |
| **0.5 (SATISFACTORY)** | Satisfactory. | Satisfactory. | Satisfactory. | Watershed has been modified, and/or loading from the watershed certainly or likely surpasses natural washout. |
| **0.4** |  |  |  |  |
| **0.3 (POOR)** | Tolerable. | Tolerable. | Tolerable. |  |
| **0.2** |  |  |  |  |
| **0.1 (VERY POOR)** | Poor. | Poor. | Poor. | Watershed is mostly constructed/modified (e.g. population centres, agriculture), and/or loading from the watershed is certainly or likely very pronounced, transgressing natural washout. |
| **0.0 (Not a habitat type)** |  | | | |

#### 3.2. Ponds and lakes for which the ecological status classification of the EU Water Framework Directive (WFD) is not applicable

|  | **(Primary)**  **Aquatic and shore vegetation representativeness (vascular plants and/or lichens and/or mosses): community and vegetation structure (abundance, coverage, species/species complex abundance ratios)** | **(Primary)**  **Watershed condition** | **(Primary)**  **Water level and fluctuation** | **(Primary)**  **Overgrowth** | **(Primary for ponds, secondary for lakes)**  **Impact of factors altering the naturalness of the shore zone and adjacent zone (e.g. forest or peatland ditches, riparian forest logging, ploughing or trenching of sites)**  *Shore zone and adjacent zone: area directly impacting waterbody* | | **(Primary)**  **Invasive alien** **species** | **(Secondary)**  **Aquatic construction and benthic modification (e.g. dredging, piling)** |
| --- | --- | --- | --- | --- | --- | --- | --- | --- |
| **Relative attribute weight** | **2** | **2** | **2** | **2** | **Ponds 2** | **Lakes 1** | **2** | **1** |
| **1.0 (EXCELLENT)** | Representative vegetation. Community and vegetation structure are characteristic of habitat type, geographical area, and local conditions. No signs of altered community or community structure, e.g. due to eutrophication. | Watershed is pristine or near-natural, and loading from watershed does not surpass natural washout (e.g. nutrients, solid particles, humus). | Water level is natural. and water level fluctuation is characteristic of habitat type and local conditions and is undisturbed. | Overgrowth or overgrowth acceleration due to anthropogenic activity are not present. | Shore zone and adjacent zone are pristine or near-natural in characteristic manner of habitat type and local conditions, no signs of anthropogenic activity. | | No invasive alien species. | Benthic shows no signs of aquatic construction or modification. |
| **0.9** |  |  |  |  |  | |  |  |
| **0.8** |  |  |  |  |  | |  |  |
| **0.7 (GOOD)** |  |  |  | Signs of overgrowth are present, but overgrowth is not particularly rapid (verifiable e.g. using aerial photographs), or overgrowth likely does not threaten the entire benthic habitat e.g. due to water depth. | Shore zone and/or adjacent zone are lightly modified. | |  | Benthic shows signs of construction or modification, but aquatic vegetation characteristic of habitat type has recovered. |
| **0.6** |  |  |  |  |  | |  |  |
| **0.5 (SATIS-FACTORY)** | Vegetation representativeness has decreased. Some species characteristic of habitat type are present, but community and/or its structure are altered. | Watershed has been modified, and/or loading from the watershed certainly or likely surpasses natural washout. | Clear changes in water level and/or water level variability due e.g. to waterbody regulation. |  | Shore zone and/or adjacent zone have bee moderately modified. | | Invasive alien species are present, but currently not at levels that impede the structure or functioning of the habitat type. However, a distinct risk exists that levels will increase to impeding. | Benthic has been constructed or modified, but aquatic vegetation characteristic of habitat type shows signs of recovery. |
| **0.4** |  |  |  |  |  | |  |  |
| **0.3 (POOR)** |  |  |  | Clear overgrowth, overgrowth is rapid (verifiable e.g. using aerial photographs), complete benthic overgrowth is possible. | Shore zone and/or adjacent zone have been greatly modified. | |  |  |
| **0.2** |  |  |  |  |  | |  |  |
| **0.1 (VERY POOR)** | Vegetation is unrepresentative. Community is greatly altered, and/or species characteristic of habitat type and site are barely present at all, vegetation structure is uncharacteristic of habitat type. | Watershed is mostly constructed/modified (e.g. population centres, agriculture), and/or loading from the watershed is certainly or likely very pronounced, transgressing natural washout. | Water level is greatly altered, and/or water level variability is non-natural e.g. due to waterbody regulation. | Fully overgrown. | Shore zone and adjacent zone have been greatly modified. | | Structure and/or functioning of habitat type is clearly reduced due to invasive alien species. | Benthic has been greatly modified (e.g. dredged, blasted) or is completely constructed, coated, or artificial and does not uphold community characteristic of habitat type, community cannot recover in current conditions. Pronounced erosion or extensive water traffic may also be present (e.g. a harbour). |
| **0.0 (Not a habitat type)** |  | | | | | | | |

### 4. Inland waters and shores: spring complexes

#### 4.1. All spring habitat types

|  | **Primary:**  **Spring-impacted surface area (including bog pools, quagmires, seepages etc. and the complexes formed by these)** | **Secondary:**  **Anthropogenic activity influencing spring complex (e.g. weeding, ditching, water extraction structures such as digging, well rings, paving)** | **Secondary: Yield** | **Secondary:**  **Location of ground water seepage** | **Secondary:**  **Surface water effect and water quality** | **Secondary:**  **Anthropogenic activity that impacts adjacent environment (e.g. earth extraction, arable land clearance, road network, forestry)**  *Adjacent environment: area that has direct impact on spring complex* |
| --- | --- | --- | --- | --- | --- | --- |
| **Relative attribute weight** | **2** | **1** | **1** | **1** | **1** | **1** |
| **1.0 (EXCELLENT)** | No signs that anthropogenic activity has decreased surface area of spring complex. | Spring complex is pristine or near-natural, no signs of anthropogenic activity. | Seepage shows no signs of decline due to anthropogenic activity. | Ground water is seeping through only in natural spots. | No clear surface water effect is apparent, and water quality is pristine or near-natural, as characteristic of habitat type and local conditions. | Shore and adjacent environment are pristine or near-natural, as characteristic of habitat type and local conditions, no signs of anthropogenic activity. |
| **0.9** |  |  |  |  |  |  |
| **0.8** |  |  |  |  |  |  |
| **0.7 (GOOD)** | Original natural spring-impacted surface area has decreased. Spring impact is present not only in pristine parts but also in areas of spring complex altered by anthropogenic activity. | Structural attributes of spring complex are slightly weakened due to anthropogenic activity. |  |  |  | Shore zone and adjacent zone have been slightly modified. |
| **0.6** |  |  |  |  |  |  |
| **0.5 (SATISFACTORY)** |  |  |  | Ground water is seeping through in natural spots, but seepage also occurs elsewhere in adjacent environment due to anthropogenic activity. |  | Shore zone and adjacent zone have been moderately modified. |
| **0.4** |  |  |  |  |  |  |
| **0.3 (POOR)** | Original natural spring-impacted surface area is markedly decreased due to anthropogenic activity. | Structucal characteristics of spring complex are greatly decreased due to anthropogenic activity. |  |  |  | Shore zone and adjacent zone have been greatly modified. |
| **0.2** |  |  |  |  |  |  |
| **0.1 (VERY POOR)** | No or very little spring-impacted surface area remaining. | Structural attributes of spring complex are intensely weakened due to anthropogenic activity. | Spring yield is greatly reduced or has ceased due to anthropogenic activity. | Ground water no longer seeping through in natural spots but only elsewhere in the adjacent environment due to anthropogenic activity. | Surface water flow into site is marked, and water quality has consequently decreased or is altered (e.g. water brownification, cloudiness). | Shore zone and adjacent zone have been intensely modified. |
| **0.0 (Not a habitat type)** |  | | | | | |

### 5. Inland waters and shores: running waters

#### 5.1. All running waters

|  | **(Primary)**  **Structural attributes of channel are characteristic of habitat type and local conditions (e.g. varying grain size, rapids, pools, winding, meandering) and change with anthropogenic activity (e.g. bushing, straightening, construction)** | **(Primary)**  **Watershed condition** | **(Secondary)**  **Vegetation representativeness (vascular plants and/or mosses): community and vegetation structure (abundance, coverage, species/species complex abundance ratios)** | **(Secondary)**  **Regulation** | **(Secondary)**  **Accessibility (dams, culverts, and other barriers** | **(Secondary)**  **Impact of anthropogenic activity that modified shore zone and adjacent zone (e.g. ditching, riparian forest cuttings, ploughing or trenching of logging sites, construction)**  *Shore and adjacent zone: area that has direct impact on running water* |
| --- | --- | --- | --- | --- | --- | --- |
| **Relative attribute weight** | **2** | **2** | **1** | **1** | **1** | **1** |
| **1.0 (EXCELLENT)** | Structural attributes are pristine or near-natural, no signs of anthropogenic activity. | Watershed is pristine or near-natural, and loading from watershed does not surpass natural washout (e.g. nutrients, solid particles, humus). | Representative vegetation. Community and vegetation structure are characteristic of habitat type, geographical area, and local conditions. | Water level fluctuation is chacteristic of habitat type and local conditions and is undisturbed. | No man-made barriers in channel. | Shore zone and adjacent zone are pristine or near-natural in characteristic manner of habitat type and local conditions, no signs of anthropogenic activity. |
| **0.9** |  |  |  |  |  |  |
| **0.8** |  |  |  |  |  |  |
| **0.7 (GOOD)** | Structural attributes are slightly weakened due to anthropogenic activity. |  |  |  |  | Shore zone and/or adjacent zone have been lightly modified. |
| **0.6** |  |  |  |  |  |  |
| **0.5 (SATISFACTORY)** | Structural attributes are moderately weakened due to anthropogenic activity. | Watershed has been modified, and/or loading from the watershed certainly or likely surpasses natural washout. | Vegetation representativeness is decreased. Some species characteristic of habitat type are present, but community and/or its structure are altered. | Water level fluctuation shows clear changes, for example due to waterbody regulation. | Barriers and/or impediments in channel (e.g. drums) may hinder the movements of certain organisms. | Shore zone and/or adjacent zone have been moderately modified. |
| **0.4** |  |  |  |  |  |  |
| **0.3 (POOR)** | Structucal attributes are greatly weakened due to anthropogenic activity. |  |  |  | Barriers in channel (e.g. dams with technical fishways) hinder the movements of most organisms up- or downstream. | Shore zone and/or adjacent zone have been greatly modified. |
| **0.2** |  |  |  |  |  |  |
| **0.1 (VERY POOR)** | Completely modified channel section. | Watershed is mostly constructed/modified (e.g. population centres, agriculture), and/or loading from the watershed is certainly or likely very pronounced, transgressing natural washout. | Vegetation is unrepresentative. Community is greatly altered, and/or species characteristic of habitat type and site are barely present at all, vegetation structure is uncharacteristic of habitat type. | Water level fluctuation is not natural, e.g. due to waterbody regulation. | Channel has man-made barrier(s) (e.g. dams) that markedly prevent or completely inhibit organisms from moving up- or downstream. | Shore zone and/or adjacent zone have been very intensely modified. |
| **0.0 (Not a habitat type)** |  | | | | | |

### 6. Inland water and shores: Shore habitat types of inland waters

#### 6.1. All shore habitat types of inland waters

|  | **(Primary)**  **Vegetation representativeness (vascular plants and/or lichens and/or mosses): community and vegetation structure (abundance, coverage, species/species complex abundance ratios)** | **(Primary)**  **Water level fluctuation** | **(Primary)**  **Construction, modification, other anthropogenic impact** | **(Secondary)**  **Invasive alien** **plant species** |
| --- | --- | --- | --- | --- |
| **Relative attribute weight** | **2** | **2** | **2** | **1** |
| **1.0 (EXCELLENT)** | Representative vegetation. Community and vegetation structure are characteristic and representative of habitat type, geographical area, local conditions (including shore zone width and topography). | Water level fluctuation is natural and undistuturbed or comparable. Vegetation community may show zoning in a manner characteristic of habitat type and local conditions. | No signs or very few signs of anthropogenic activity. Shore is completely unconstructed. At most, individual paths or duckboards and light recreational use may be present. | No invasive alien plant species present. |
| **0.9** |  |  |  |  |
| **0.8** |  |  |  |  |
| **0.7 (GOOD)** | Vegetation representativeness is decreased to a degree. Community characteristic of habitat type is observable but is not as representative as in the ‘Excellent’ category. Vegetation structure may be altered to a degree. May show small signs of overgrowth or erosion. |  | Minor anthropogenic activity. Small-scale construction and/or modification in a way that barely affects habitat type (paths, docks, small boat ramp, light erosion). |  |
| **0.6** |  |  |  |  |
| **0.5 (SATISFACTORY)** | Vegetation representativeness is distinctly decreased. Community and/or vegetation structure exhibit clear changes, some representative species are present. May show clear signs of overgrowth or erosion. | Water level fluctuation shows clear changes e.g. due to waterbody regulation. |  | Individual invasive alien plants present. |
| **0.4** |  |  |  |  |
| **0.3 (POOR)** |  |  | Fairly abundant anthropogenic activity. Shore may be constructed, modified, or eroded in a way tht impacts habitat type and/or its rehabilitation alternatives (e.g. numerous recreational cabins, dredging, regular traffic, erosion). | Several occurrences of invasive alien plants. |
| **0.2** |  |  |  |  |
| **0.1 (VERY POOR)** | Vegetation is unrepresentative. Community is greatly altered, and/or species characteristic of habitat type and site are barely present at all, vegetation structure is uncharacteristic of habitat type.  May be extensively overgrown or eroded. | Water level fluctuation is not natural at all e.g. due to waterbody regulation. | Intensive anthropogenic activity. Shore may be constructed, modifiied, extensively eroded, or artificial througout. | Area largerly taken over by invasive alien plants. |
| **0.0 (Not a habitat type)** |  | | | |

### 7. Peatlands

#### 7.1. Open peatlands

|  | **(Primary)**  **Representativeness of peatland vegetation (vascular plants and/or mosses): community and vegetation structure (abundance, coverage, species/species complex abundance ratios)** | **(Primary)**  **Hydrology** | **(Primary)**  **Peatland’s relationship with environment (condition of peatland complex)**  *Not used on boreal mire complexes or on individual peatland patches consisting of one peatland type.* | **(Primary)**  **Overgrowth** | **(Secondary)**  **Other anthropogenic impact** |
| --- | --- | --- | --- | --- | --- |
| **Relative attribute weight** | **2** | **2** | **2** | **2** | **1** |
| **1.0 (EXCELLENT)** | Representative vegetation. Community and vegetation structure are characteristic and representative of habitat type, geographical area, local conditions, and natural successional stage. | Pristine hydrology characteristic of habitat type. | Open areas of peatland complex are pristine or near-natural. | No overgrowth. At most, individual trees/scrubs, as characteristic of the habitat type. | No signs or very few signs of anthropogenic activity. At most, individual paths, duckboards, birdwatching towers are possible. No harmful alien plants. |
| **0.9** |  |  |  |  |  |
| **0.8** |  |  |  |  |  |
| **0.7 (GOOD)** | Vegetation representativeness is decreased to a degree. Essential community characteristic of habitat type is observable but is not as representative as in the ‘excellent’ category. Vegetation structure may be altered to a degree. | Small signs of hydrological alteration either due to ditching or surrounding land use. |  | Traces of tree sprouting, shrub encroachment, or reed encroachment. | Minor anthropogenic activity. For example, paths, tracks, slight terrain erosion. Individual invasive alien plants may be present. |
| **0.6** |  |  |  |  |  |
| **0.5 (SATISFACTORY)** |  | Clear hydrological changes either due to ditching or surrounding land use. | Open areas of peatland complex have partially weakened (e.g. partial ditching). |  | Fairly abundant anthropogenic activity. For example, paths, tracks, clear terrain erosion, and/or anthropogenic impact related to light, noise, or air quality. Moderate levels of invasive alien plants may be present. |
| **0.4** |  |  |  |  |  |
| **0.3 (POOR)** | Vegetation representativeness is distinctly decreased. Community and/or vegetation structure exhibit clear changes, some representative species are present. |  |  | Clear tree sprouting, shrub encroachment, or reed encroachment, individual trunk trees may be present. |  |
| **0.2** |  |  |  |  |  |
| **0.1 (VERY POOR)** | Vegetation is unrepresentative. Community is greatly altered, and/or species characteristic of habitat type and site are barely present, vegetation structure uncharacteristic of habitat type. | Hydrology is greatly altered due to ditching or surrounding land use. | Peatland complex is weakened throughout (e.g. ditching throughout). | Pronounced overgrowth. Abundant trunk trees, possible abundant seedlings/saplings and scrub, or pronounced reed encroachment. | Intensive anthropogenic activity. For example, marked terrain erosion, built structures. Invasive alien plants may be present in abundance. |
| **0.0 (Not a habitat type)** |  | | | | |

#### 7.2. Wooded peatlands

|  | **(Primary)**  **Peatland vegetation representativeness (vascular plants and/or lichens and/or mosses): community and vegetation structure (abundance, coverage, species/species complex abundance ratios)** | **(Primary)**  **Hydrology** | **(Primary)**  **Peatland’s relationship with environment (condition of peatland complex)**  *Not used on boreal mire complexes or on individual peatland patches consisting of one peatland type.* | **(Primary)**  **Forest structure (tree density, age distribution, spatial distribution, crown stratification, species distribution)** | **(Secondary)**  **Dead wood quantity** | **(Secondary)**  **Other anthropogenic impact** |
| --- | --- | --- | --- | --- | --- | --- |
| **Relative attribute weight** | **2** | **2** | **2** | **2** | **1** | **1** |
| **1.0 (EXCELLENT)** | Representative vegetation. Community and vegetation structure are characteristic and representative of habitat type, geographical area, local conditions, and natural successional stage. | Pristine hydrology characteristic of habitat type. | Forested areas of peatland complex are pristine or near-natural throughout. | All structural attributes characteristic of habitat type, geographical area, local conditions, and natural successional stage are present. | Typical amount of pristine conditions characteristic of habitat type, geographical area, local conditions, and natural successional stage. | No signs or very few signs of anthropogenic activity. At most, individual paths, duckboards, birdwatching towers are possible. No invasive alien plants. |
| **0.9** |  |  |  |  |  |  |
| **0.8** |  |  |  |  |  |  |
| **0.7 (GOOD)** | Vegetation representativeness is decreased to a degree. Essential community characteristic of habitat type is observable but is not as representative as in the ‘excellent’ category. Vegetation structure may be altered to a degree. | Small signs of hydrological alteration either due to ditching or surrounding land use. |  | Forest structure is slightly altered, e.g. due to management or tree sprouting caused by desiccation. |  | Minor anthropogenic activity. For example, paths, tracks, slight terrain erosion. Low levels of invasive alien plants may occur. |
| **0.6** |  |  |  |  |  |  |
| **0.5 (SATISFACTORY)** |  | Clear hydrological changes either due to ditching or surrounding land use. | Forested areas of peatland complex are partially weakened (e.g. partial ditching or logging). |  | Less dead wood present than would be characteristic for habitat type and local conditions. | Fairly abundant anthropogenic activity. For example, trails, tracks, clear terrain erosion. Moderate levels of invasive alien plants may occur. |
| **0.4** |  |  |  |  |  |  |
| **0.3 (POOR)** | Vegetation representativeness is distinctly decreased. Community and/or vegetation structure exhibit clear changes, some representative species are present. |  |  | Forest structure is clearly altered, e.g. due to management or tree sprouting caused by desiccation. |  |  |
| **0.2** |  |  |  |  |  |  |
| **0.1 (VERY POOR)** | Vegetation is unrepresentative. Community is greatly altered, and/or species characteristic of habitat type and site are barely present, vegetation structure is uncharacteristic of habitat type. | Hydrology is greatly altered due to ditching or surrounding land use. | Forested areas of peatland complex are weakened throughout (e.g. ditching or logging throughout). | Forest structure is uncharacteristic of habitat type. | No dead wood or nearly no dead wood present. | Intensive anthropogenic activity. For example, marked terrain erosion, built structures. Invasive alien plants may be present in abundance. |
| **0.0 (Not a habitat type)** |  | | | | | |

### 8. Forests

#### 8.1. Herb-rich heath forests, Hardwood forests on podsolic soils, Mesic heath forests, Sub-xeric heath forests, Xeric heath forests, and Herb-rich, Vaccinium myrtillus, Vaccinium vitis-idaea, and dwarf-shrub drained peatland forests

|  | **(Primary)**  **Development class** | **(Primary)**  **Occurrence of structural tree attributes characteristic of habitat type: uneven-agedness, crown stratification, stochastic spatial distribution, fire scars, tree species richness** | **(Primary)**  **Dead wood quantity^3^ and structural attributes characteristic of habitat type: continuum, large-diameter dead wood, tree species richness**  *Large-diameter dead wood:*  *basal diameter ≥30 cm* | **(Secondary)**  **Vegetation representativeness (vascular plants and/or lichens and/or mosses): community and vegetation structure (abundance, coverage, species/species complex abundance ratios)** | **(Secondary)**  **Number of large-diameter trees^4^**  *A large-diameter living tree: diameter at breast height ≥ 40 cm* | **(Secondary)**  **Invasive alien** **plant species** | **(Secondary)**  **Other anthropogenic impact** |
| --- | --- | --- | --- | --- | --- | --- | --- |
| **Relative attribute weight** | **2** | **2** | **2** | **1** | **1** | **1** | **1** |
| **1.0 (EXCELLENT)** | Old forest^1^ or naturally generated forest of a previous successional stage (e.g. after land uplift, fire, or storm). | All structural attributes typical of habitat type, local conditions, and natural successional stage. | Naturally generated dead wood characteristic of habitat type. All dead wood structural attributes characteristic of habitat type are present. | Representative vegetation. Community and vegetation structure are characteristic and representative of habitat type, geographical area, local conditions, and natural successional stage. | Quantity characteristic of habitat type, local conditions, and natural successional stage. | No invasive alien plant species present. | No signs or very few signs of anthropogenic activity. For example, individual paths or forest grazing are possible. |
| **0.9** |  |  |  |  |  |  |  |
| **0.8** |  |  |  |  |  |  |  |
| **0.7 (GOOD)** | Mature commercial forest stand or older. | At least three structural attributes present. | At least moderate quantities of naturally generated dead wood and at least two structural attributes present. | Vegetation representativeness is decreased to a degree. Essential community characteristic of habitat type is observable but is not as representative as in the ‘excellent’ category. Vegetation structure may be altered to a degree. | Large-diameter trees occur but in numbers smaller than characteristic of habitat type, local conditions, and natural successional stage. |  | Minor anthropogenic activity. For example, harvester tracks or individual blocked forest ditches. |
| **0.6** |  |  |  |  |  |  |  |
| **0.5 (SATISFACTORY)** | Young–middle-aged commercial forest or shelterwood stand or seed-tree stand^2^. | Two structural attributes. | Low levels of naturally generated dead wood or moderate levels of artificially produced dead wood, and at least one structural attribute is present. | Vegetation representativeness is distinctly decreased. Community and/or vegetation structure exhibit clear changes, some representative species are present. | Individual large-diameter trees. | Individual invasive alien plants present. | Moderate anthropogenic activity. For example, terrain erosion, littering, or forest ditches. |
| **0.4** |  |  |  |  |  |  |  |
| **0.3 (POOR)** | Seedling or sapling stand or seed-tree stand^2^. | One structural attribute, or non-native mixed-species forest. | Individual dead wood objects that are not large in diameter, no stuctural attributes are present. |  |  | Several occurrences of invasive alien plants. |  |
| **0.2** |  |  |  |  |  |  |  |
| **0.1 (VERY POOR)** | Cleart-cut area or seed-tree stand^2^. | No structural attributes are present, or a single-species non-native forest. | No dead wood present. | Vegetation is unrepresentative. Community is greatly altered, and/or species characteristic of habitat type and site are barely present, vegetation structure is uncharacteristic of habitat type. | No large-diameter trees present. | Area largerly taken over by invasive alien plants. | Intensive anthropogenic activity. For example, marked terrain erosion, wide-scale forest ditching, soil tillage, or significant littering. |

^1^Age limits (in years) according to most compatible heath forest type description in the Finnish Red List of Habitat Types. Heath forest reference values are applied to analogous drained peatland forest.

|  | Xeric heath forest | Sub-xeric heath forest | Mesic heath forest, coniferous | Mesic heath forest, deciduous | Herb-rich heath forest and herb-rich forest, coniferous | Herb-rich heath forest and herb-rich forest, deciduous |
| --- | --- | --- | --- | --- | --- | --- |
| Southern Finland | 160 | 140 | 140 | 80 | 120 | 80 |
| North Ostrobothnia and Kainuu | 200 | 160 | 140 | 80 | 140 | 80 |
| Koillismaa and southern Lapland | 200 | 180 | 160 | 80 | 160 | 80 |
| Mid-Lapland | 200 | 200 | 200 | 80 | 180 | 80 |
| Upper Lapland and protection forest area | 220 | 200 | 200 | 100 | 200 | 100 |

^2^Seed-tree stand status is defined based on lower tree stratum, i.e. seedling material.

^3^Dead wood quantity categories. Natural quantity in particular is indicative. Heath forest reference values are applied to analogous drained peatland forest.

|  |  | Mesic and herb-rich heath forests | Xeric and sub-xeric heath forests |
| --- | --- | --- | --- |
| Dead wood, natural quantity (m^3^/ha) | Southern Finland | 30 | 20 |
|  | Northern Finland | 20 | 10 |
| Dead wood, moderate quantity (m^3^/ha) | Southern Finland | 10 | 5 |
|  | Northern Finland | 5 | 3 |
| Dead wood, small quantity (m^3^/ha) | Southern Finland | 5 | 2 |
|  | Northern Finland | 2 | No indicative lower bound |

^4^Quantity categories for pristine levels of large-diameter trees (quantities may be smaller in northern Finland):

- Herb-rich heath forests, mesic heath forests, sub-xeric heath forests, and hardwood forests on podsolic soils, herb-rich, herb-rich heath, *Vaccinium myrtillus,* and *Vaccinium vitis-idaea* drained peatlands: 20 trunks/ha.
- Xeric heath forests, dwarf-shrub drained peatland forests: 10 trunks/ha.

#### 8.2. Herb-rich heath forests, Herb-rich forests with broadleaved deciduous trees, Esker forests, Inland dune forests, Forests on ultrabasic soils

|  | **(Primary)**  **Development class** | **(Primary)**  **Occurrence of structural tree attributes characteristic of habitat type: uneven-agedness, crown stratification, stochastic spatial distribution, fire scars, tree species richness** | **(Primary)**  **Dead wood quantity^3^ and structural attributes characteristic of habitat type: continuum, large-diameter dead wood, tree species richness**  *Large-diameter dead wood:*  *basal diameter of ≥ 30 cm* | **(Primary) Vegetation representativeness (vascular plants and/or lichens and/or mosses): community and vegetation structure (abundance, coverage, species/species complex abundance ratios)** | **(Secondary)**  **Number of large-diameter trees^4^**  *A large-diameter living tree: diameter at breast height ≥ 40 cm* | **(Secondary)**  **Invasive alien** **plant species** | **(Secondary)**  **Other anthropogenic impact** | **(Secondary)**  **Additionally for moist herb-rich forests: hydrology** |
| --- | --- | --- | --- | --- | --- | --- | --- | --- |
| **Relative attribute weight** | **2** | **2** | **2** | **2** | **1** | **1** | **1** | **Moist herb-rich forests 1, not applicable to other habitat types** |
| **1.0 (EXCELLENT)** | Old forest^1^ or naturally generated forest of a previous successional stage (e.g. after land uplift, fire, or storm). | All structural attributes typical of habitat type, local conditions, and natural successional stage. | Naturally generated dead wood in pristine or near-natural quantities. All dead wood structural attributes characteristic of habitat type are present. | Representative vegetation. Community and vegetation structure are characteristic and representative of habitat type, geographical area, local conditions, and natural successional stage. | Quantity characteristic of habitat type, local conditions, and natural successional stage. | No invasive alien plant species present. | No signs or very few signs of anthropogenic activity. For example, individual paths or forest grazing are possible | Natural or near-natural, stable hydrology. |
| **0.9** |  |  |  |  |  |  |  |  |
| **0.8** |  |  |  |  |  |  |  |  |
| **0.7 (GOOD)** | Mature commercial forest stand or older. | At least three structural attributes present. | At least moderate quantities of naturally generated dead wood and at least two structural attributes present. | Vegetation representativeness is decreased to a degree. Essential community characteristic of habitat type is observable but is not as representative as in the ‘excellent’ category. Vegetation structure may be altered to a degree. | Large-diameter trees occur but in numbers smaller than characteristic of habitat type, local conditions, and natural successional stage. |  | Minor anthropogenic activity. For example, harvester tracks or individual blocked forest ditches. |  |
| **0.6** |  |  |  |  |  |  |  |  |
| **0.5 (SATISFACTORY)** | Young–middle-aged commercial forest or shelterwood stand or seed-tree stand^2^. | Two structural attributes. | Low levels of naturally generated dead wood or moderate levels of artificially produced dead wood, and at least one structural attribute present. | Vegetation representativeness is distinctly decreased. Community and/or vegetation structure exhibit clear changes, some representative species are present. | Individual large-diameter trees. | Individual invasive alien plants. | Moderate anthropogenic activity. For example, terrain erosion, littering, or forest ditches. | Hydrology is disturbed to a degree, signs of desiccation present. |
| **0.4** |  |  |  |  |  |  |  |  |
| **0.3 (POOR)** | Seedling or sapling stand or seed-tree stand^2^. | One structural attribute, or non-native mixed-species forest. | Individual dead wood objects that are not large in diameter occur, no stuctural attributes are present. |  |  | Several occurrences of invasive alien plants. |  |  |
| **0.2** |  |  |  |  |  |  |  |  |
| **0.1 (VERY POOR)** | Cleart-cut area or seed-tree stand^2^. | No structural attributes are present, or a single-species non-native forest stand. | No dead wood present. | Vegetation is unrepresentative. Community is greatly altered and/or species characteristic of habitat type and site are barely present, vegetation structure is uncharacteristic of habitat type. | No large-diameter trees present. | Area largerly taken over by invasive alien plants. | Intensive anthropogenic activity. For example, marked terrain erosion, wide-scale forest ditching, soil tillage, or significant littering. | Hydrology is greatly disturbed, clearly desiccated. |
| **0.0 (Not a habitat type)** |  | | | | | | | |

^1^Age limits (in years) according to most compatible heath forest type description in the Finnish Red List of Habitat Types. Heath forest reference values are applied to analogous drained peatland forest.

|  | Xeric heath forest | Sub-xeric heath forest | Mesic heath forest, coniferous | Mesic heath forest, deciduous | Herb-rich heath forest and  herb-rich forest, coniferous | Herb-rich heath forest and  herb-rich forest, deciduous |
| --- | --- | --- | --- | --- | --- | --- |
| Southern Finland | 160 | 140 | 140 | 80 | 120 | 80 |
| North Ostrobothnia and Kainuu | 200 | 160 | 140 | 80 | 140 | 80 |
| Koillismaa and southern Lapland | 200 | 180 | 160 | 80 | 160 | 80 |
| Mid-Lapland | 200 | 200 | 200 | 80 | 180 | 80 |
| Upper Lapland and protection forest | 220 | 200 | 200 | 100 | 200 | 100 |

^2^Seed-tree stand status isdefined based on lower tree stratum i.e. seedling material.

^3^Dead wood quantity categories. Natural quantity in particular is indicative. Heath forest reference values are applied to analogous drained peatland forest.

|  |  | Herb-rich forests | Esker forests,  inland dune forests,  forests on ultrabasic soils |
| --- | --- | --- | --- |
| Dead wood, intrinsic quantity (m^3^/ha) | Southern Finland | 30 | 20 |
|  | Northern Finland | 20 | 10 |
| Dead wood, moderate quantity (m3/ha) | Southern Finland | 10 | 5 |
|  | Northern Finland | 5 | 3 |
| Dead wood, small quantity (m^3^/ha) | Southern Finland | 5 | 2 |
|  | Northern Finland | 2 | No indicative lower bound |

^4^Quantity categories for pristine levels of large-diameter trees (Quantities may be smaller in northern Finland):

- Herb-rich forests: 30 trunks/ha
- Esker forests and inland dune forests: 10 trunks/ha (spruces are not considered)
- Forests on ultrabasic soils: 10 trunks/ha

#### 8.3. Barren heath forests, Forests on rocky terrain, *Cladonia* drained peatland forests

|  | **(Primary)**  **Development class** | **(Primary)**  **Occurrence of structural tree attributes characteristic of habitat type: uneven-agedness, crown stratification, stochastic spatial distribution, fire scars. Additionally for forests on rocky terrain: tree species richness.** | **(Primary)**  **Dead wood quantity and structural attributes characteristic of habitat type: continuum, large-diameter dead wood, multiple tree species**  Large-diameter dead wood:  basal diameter *≥* 30 cm | **(Secondary)**  **Vegetation representativeness (vascular plants and/or lichens and/or mosses): community and vegetation structure (abundance, coverage, species/species complex abundance ratios)** | **(Secondary)**  **Number of large-diameter trees^4^**  *A large-diameter living tree: diameter at breast height ≥ 30 cm* | **(Secondary)**  **Invasive alien** **plant species** | **(Secondary)**  **Other anthropogenic impact** |
| --- | --- | --- | --- | --- | --- | --- | --- |
| **Relative attribute weight** | **2** | **2** | **2** | **1** | **1** | **1** | **1** |
| **1.0 (EXCELLENT)** | Old forest or previous successional stage forest generated naturally (e.g. after land uplift, fire, or storm). No signs of forest management. | All structural attributes typical of habitat type, local conditions, and natural successional stage (note: tree species richness not required for barren heath forests or *Cladonia* drained peatland forests). | Characteristic dead wood quantity is present for habitat type, local conditions, and natural successional stage. Natural dead wood generation is undisturbed. At least one sturctural attribute is present. | Representative vegetation. Community and vegetation structure are characteristic and representative of habitat type, geographical area, local conditions, and natural successional stage. | At least individual large-diameter trees present. | No invasive alien plant species present. | No signs or very few signs of anthropogenic activity. For example, individual paths or forest grazing are possible. |
| **0.9** |  |  |  |  |  |  |  |
| **0.8** |  |  |  |  |  |  |  |
| **0.7 (GOOD)** |  | Barren heath forests and *Cladonia* drained peatland forests: at least two structural attributes.  Forests on rocky terrain: at least three structural attributes. | Dead wood is present but at lower levels than characteristic of habitat type, local conditions, and natural successional stage, and/or natural dead wood generations is slightly decreased at most. | Vegetation representativeness is decreased to a degree. Essential community characteristic of habitat type is observable but is not as representative as in the ‘excellent’ category. Vegetation structure may be altered to a degree. | No large-diameter trees present, but natural generation of large-diameter trees is not noticeably disturbed due to anthropogenic activity (e.g. forest management). |  | Minor anthropogenic activity. For example, harvester tracks or individual blocked forest ditches. |
| **0.6** |  |  |  |  |  |  |  |
| **0.5 (SATISFACTORY)** | Forest management history is apparent, but previous management actions occurred several decades ago. | Barren heath forests and *Cladonia* drained peatland forests: one structural attribute.  Forests on rocky terrain: at least two structural attributes. |  | Vegetation representativeness is distinctly decreased. Community and/or vegetation structure exhibit clear changes, some representative species are present. |  | Individual invasive alien plants. | Moderate anthropogenic activity. For example, terrain erosion, littering, or forest ditches. |
| **0.4** |  |  |  |  |  |  |  |
| **0.3 (POOR)** |  | Forests on rocky terrain: one structural attribute. | Dead wood is sparsely spaced out or occurs as individual objects, and/or natural dead wood generation is clearly decreased. |  |  | Several occurrences of invasive alien plants. |  |
| **0.2** |  |  |  |  |  |  |  |
| **0.1 (VERY POOR)** | Recently treated forest. Clear-cut area, seedlings/saplings or comparable. | No structural attributes. | No dead wood, and/or natural dead wood generation is greatly decreased. | Unrepresentative vegetation. Community is greatly altered, and/or species characteristic of habitat type and site are barely present vegetation structure is uncharacteristic of habitat type. | No large-diameter trees present, and their natural generation is clearly disturbed due to anthropogenic activity (e.g. forest management). | Area largerly taken over by invasive alien plants. | Intensive anthropogenic activity. For example, marked terrain erosion, wide-scale forest ditching, soil tillage, or significant littering. |
| **0.0 (Not a habitat type)** |  | | | | | | |

#### 8.4. Inland flooded forests

|  | **(Primary)**  **Development class** | **(Primary)**  **Occurrence of structural tree attributes characteristic of habitat type: uneven-agedness, crown stratification, stochastic spatial distribution, fire scars, tree species richness** | **(Primary)**  **Dead wood quantity and structural attributes characteristic of habitat type: continuum, large-diameter dead wood, tree species richness**  *Large-diameter dead wood:*  *basal diameter ≥ 30 cm* | **(Primary)**  **Flood conditions** | **(Secondary) Vegetation representativeness (vascular plants and/or lichens and/or mosses): community and vegetation structure (abundance, coverage, species/species complex abundance ratios)** | **(Secondary)**  **Number of large-diameter trees**  *A large-diameter living tree: diameter at breast height : ≥ 30 cm* | **(Secondary)**  **Invasive alien** **plant species** | **(Secondary)**  **Other anthropogenic impact** |
| --- | --- | --- | --- | --- | --- | --- | --- | --- |
| **Relative attribute weight** | **2** | **2** | **2** | **2** | **1** | **1** | **1** | **1** |
| **1.0 (EXCELLENT)** | Old forest or previous successional stage forest generated naturally (e.g. after land uplift, fire, or storm). No signs of forest management. | All structural attributes characteristic of habitat type, local conditions, and natural successional stage are present. | Dead wood quantity^2^ is characteristic of habitat type, local conditions, and natural successional stage, and natural dead wood generation is undisturbed, and all structural dead wood attributes are present. | Natural flood conditions or comparable. | Representative vegetation. Community and vegetation structure are characteristic and representative of habitat type, geographical area, local conditions, and natural successional stage. | At least individual large-diameter trees^.^ | No invasive alien plant species present. | No signs or very few signs of anthropogenic activity. For example, individual paths or forest grazing are possible. |
| **0.9** |  |  |  |  |  |  |  |  |
| **0.8** |  |  |  |  |  |  |  |  |
| **0.7 (GOOD)** |  | At least three structural attributes present. | Dead wood is present but at lower levels than characteristic of habitat type, local conditions, and natural successional stage, and/or natural dead wood generations is slightly decreased at most, and at least one sturctural attribute is present. |  | Vegetation representativeness is decreased to a degree. Essential community characteristic of habitat type is observable but is not as representative as in the ‘excellent’ category. Vegetation structure may be altered to a degree. | No large-diameter trees are present, but natural generation of large-diameter trees is not noticeably disturbed due to anthropogenic activity (e.g. forest management). |  | Minor anthropogenic activity. For example, harvester tracks or individual blocked forest ditches. |
| **0.6** |  |  |  |  |  |  |  |  |
| **0.5 (SATISFACTORY)** | Forest management history is apparent, but previous management actions occurred several decades ago. | Two structural attributes present. |  | Flood conditions moderately altered e.g. due to waterbody regulation. | Vegetation representativeness is distinctly decreased. Community and/or vegetation structure exhibit clear changes, some representative species are present. |  | Individual invasive alien plants. | Fairly abundanct anthropogenic activity. For example, terrain erosion, littering, or forest ditching. |
| **0.4** |  |  |  |  |  |  |  |  |
| **0.3 (POOR)** |  | One structural attribute present. | Dead wood os sparsely spaced out or occurs as individual objects, and/or natural dead wood generation is clearly decreased. No structural attributes. |  |  |  | Several occurrences of invasive alien plants. |  |
| **0.2** |  |  |  |  |  |  |  |  |
| **0.1 (VERY POOR)** | Recently treated forest. Clear-cut area, seedlings/saplings or comparable. | No structural attributes present. | No dead wood, and/or natural dead wood generation is greatly decreased. | Flood conditions markedly altered, e.g. due to waterbody regulation. | Vegetation is unrepresentative. Community is greatly altered and/or species characteristic of habitat type and site are barely present, vegetation structure is uncharacteristic of habitat type. | No large-diameter trees are present, and their natural generation is clearly disturbed due to anthropogenic activity (e.g. forest management). | Area largerly taken over by invasive alien plants. | Intensive anthropogenic activity For example, marked terrain erosion, wide-scale forest ditching, soil tillage, or significant littering. |
| **0.0 (Not a habitat type)** |  | | | | | | | |

### 9. Rock outcrops and scree

#### 9.1. Rock outcrops

|  | **(Primary)**  **Vegetation representativeness (vascular plants and/or lichens and/or mosses): community and vegetation structure (abundance, coverage, species/species complex abundance ratios)** | **(Primary for acidic rock outcrops, secondary for intermediate–basic, calcareous, and serpentine rock outcrops)**  **Morphological diversity of rock outcrops** | | **(Secondary)**  **Other anthropogenic impact** | **(Secondary)**  **Additionally for rock outcrops with trees: forest** | **(Primary)**  **Additionally for calcareous and serpentine rock outcrops: habitat specialist vascular plant and/or moss and/or lichen community** |
| --- | --- | --- | --- | --- | --- | --- |
| **Relative attribute weight** | **2** | **Acidic rock outcrops 2** | **Intermediate–basic rock outcrops, calcareous rock outcrops, and serpentine rock outcrops 1** | **1** | **1** | **2** |
| **1.0 (EXCELLENT)** | Representative vegetation. Community and vegetation structure are characteristic and representative of habitat type, geographical area, local conditions, and natural successional stage. | Broad rock outcrop with a variable surface profile (e.g. plateaus, vertical surfaces, crevices, tall rock faces, overhanging rock faces, caves, or cavities). | | No signs or very few signs of anthropogenic activity. At most, individual narrow paths. | Old trees and dead wood in quantities characteristic of habitat type and local conditions. | Abundant levels of habitat specialist indicator species and high species richness. |
| **0.9** |  |  | |  |  |  |
| **0.8** |  |  | |  |  |  |
| **0.7 (GOOD)** | Vegetation representativeness is decreased to a degree. Essential community characteristic of habitat type is observable but is not as representative as in the ‘excellent’ category. Vegetation structure may be altered to a degree. | Small-sized rock outcrop with variable surface profile or broad rock outcrop with a monotonic suface profile. | | Minor anthropogenic activity. Some terrain erosion possible. | Old trees are present, but no dead wood. | Moderate levels of habitat specialist species. |
| **0.6** |  |  | |  |  |  |
| **0.5 (SATISFACTORY)** |  |  | |  |  |  |
| **0.4** |  |  | |  |  |  |
| **0.3 (POOR)** | Vegetation representativeness is distinctly decreased. Community and/or vegetation structure exhibit clear changes, some representative species are present. | Small-sized rock outcrop with an only slightly variable surface profile. | | Moderate anthropogenic activity. For example, clear terrain erosion. | Only young trees present, no old trees or dead wood. | Individual habitat specialist species present. |
| **0.2** |  |  | |  |  |  |
| **0.1 (VERY POOR)** | Unrepresentative vegetation. Community is greatly altered, and/or species characteristic of habitat type and site are barely present, vegetation structure is uncharacteristic of habitat type. | Small-sized and flat rock outcrop. | | Intensive anthropogenic activity. For example, marked terrain erosion or rock outcrop quarried until bare. | Overly dense seedling/sapling stand in relation to habitat type and local conditions. | No habitat specialist species. |
| **0.0 (Not a habitat type)** |  | | | | | |

#### 9.2. Rock faces

|  | **(Primary)**  **Vegetation representativeness (vascular plants and/or lichens and/or mosses): community and vegetation structure (abundance, coverage, species/species complex abundance ratios)** | **(Primary for acidic rock faces, secondary for intermediate–basic, calcareous, and serpentine rock faces). Morphological diversity of rock face.** | | **(Primary)**  **Additionally for shady rock faces: shading and microclimate stability** | **(Secondary)**  **Additionally for rock faces with trees: trees on the rock face and at its base** | **(Primary)**  **Additionally for calcareous and serpentine rock faces:** **habitat specialist species** |
| --- | --- | --- | --- | --- | --- | --- |
| **Relative attribute weight** | **2** | **Acidic rock faces 2** | **Intermediate–basic, calcareous, and serpentine rock faces 1** | **Shady rock faces 2, not applicable to other rock face types** | **Rock faces with trees 1, not applicable to other rock face types** | **Calcareous and serpentine rock faces 2, not applicable to other rock face types** |
| **1.0 (EXCELLENT)** | Representative vegetation. Community and vegetation structure are characteristic and representative of habitat type, geographical area, local conditions, and natural successional stage. | High and broad rock face (over 10 m in height for acidic rock faces, no height limit for other rock face types). Abundant surfaces with varied moisture and other conditions (e.g. terraces, vertical surfaces, seepage water, and various shoreline-altered areas for shoreline rock faces) | | Gorge or other permanently shading structure that ensures a highly stable microclimate. | Uneven-aged forest, old trees, and dead wood present in a manner characteristic of habitat type and local conditions. | Abundant levels of habitat specialist indicator species. High species richness. |
| **0.9** |  |  | |  |  |  |
| **0.8** |  |  | |  |  |  |
| **0.7 (GOOD)** | Vegetation representativeness decreased to a degree. Essential community characteristic of habitat type is observable but is not as representative as in the ‘excellent’ category. Vegetation structure may be altered to a degree. |  | | Forest adjacent to rock face shades rock face, ensuring a moist and stable microclimate. |  | Moderate levels of habitat specialist species. |
| **0.6** |  |  | |  |  |  |
| **0.5 (SATISFACTORY)** |  | Several surfaces that vary in moisture and other conditions. | |  | Due to anthropogenic activity, only middle-aged trees present at most, with small or no quantities of dead wood. |  |
| **0.4** |  |  | |  |  |  |
| **0.3 (POOR)** | Vegetation representativeness distinctly decreased. Community and/or vegetation structure exhibit clear changes, some representative species present. |  | | Reduced shading, e.g. forest has been thinned. |  | Individual habitat specialist species. |
| **0.2** |  |  | |  |  |  |
| **0.1 (VERY POOR)** | Unrepresentative vegetation. Community is greatly altered, and/or species characteristic of habitat type and site are barely present, vegetation structure is uncharacteristic of habitat type. | Homogenous surface in terms of moisture and other conditions. | | Rock face has lost all shading (e.g. shade-producing forest has been removed), causing microclimate to become dry and extreme. | Only young trees are present due to anthropogenic activity. | No habitat specialist species. |
| **0.0 (Not a habitat type)** |  | | | | | |

#### 9.3. Scree

|  | **(Secondary for calcareous and serpentine screes, only measure for other scree types)**  **Size and variability** | **(Primary)**  **Additionally for calcareous and serpentine screes:** **habitat specialist species** |
| --- | --- | --- |
| **Relative attribute weight** | **1** | **Calcareous and serpentine screes 2: not applied to other scree types** |
| **1.0 (EXCELLENT)** | Wide scree area with varying microclimate patches (e.g. forest edge and/or shoreline, rock face bottom scree, bare inner area). Microclimate extremeness varies between scree areas. | Abundant levels of habitat specialist indicator species. High species richness. |
| **0.9** |  |  |
| **0.8** |  |  |
| **0.7 (GOOD)** | Small-sized and/or microclimatically homogenous scree, but showing bare scree surface. | Moderate levels of habitat specialist species. |
| **0.6** |  |  |
| **0.5 (SATISFACTORY)** |  |  |
| **0.4** |  |  |
| **0.3 (POOR)** | Mostly overgrown, scree surface scantily visible. | Individual habitat specialist species. |
| **0.2** |  |  |
| **0.1 (VERY POOR)** | Fully overgrown, scree surface no longer visible. | No habitat specialist species. |
| **0.0 (Not a habitat type)** |  | |

### 10. Seminatural grasslands and grazed woodlands (traditional biotopes)

#### 10.1. Traditional biotopes with an existing value category in the national traditional landscape inventory

|  | **Traditional landscape category^1^** |
| --- | --- |
| **Relative attribute weight** | **1** |
| **1.0 (EXCELLENT)** | V National |
| **0.9** | M+ regional + (close to national level) |
| **0.8** |  |
| **0.7 (GOOD)** | M regional |
| **0.6** | M­– regional - (close to local level) |
| **0.5 (SATISFACTORY)** | P+ local + (close to regional level) |
| **0.4** | L local |
| **0.3 (POOR)** | P– local– (barely exhibiting traditional landscape values) |
| **0.2** | K repairable |
| **0.1 (VERY POOR)** | No value as a traditional landscape |
| **0.0 (Not a habitat type)** |  |

^1^Kemppainen 2017. Perinnemaisemien inventointiohje. Varsinais-Suomen elinkeino-, liikenne- ja ympäristökeskuksen raportteja 25/2017. In Finnish.

#### 10.2. Open traditional biotopes

|  | **(Primary) Vegetation representativeness (vascular plants and/or lichens and/or mosses): community and vegetation structure (abundance, coverage, species/species complex abundance ratios)** | **(Primary)**  **Notable community** | **(Primary)**  **Eutrophication and problematic species** | **(Secondary)**  **Reaping or grazing that does not cause eutrophication, or other comparable management** | **(Secondary)**  **Overgrowth** | **(Secondary)**  **Invasive alien** **plant species** |
| --- | --- | --- | --- | --- | --- | --- |
| **Relative attribute weight** | **2** | **2** | **2** | **1** | **1** | **1** |
| **1.0 (EXCELLENT)** | Representative vegetation. Community and vegetation structure are charcteristic and representative of habitat type, geographical area, and local conditions. | An abundance of regionally notable traditional biotope plant species^1^ and/or a significant site for other notable or endangered biota (e.g. abundant food plants of endangered insects, or knowledge of a viable population of an endangered community). | No marked eutrophication or problematic species. | Ongoing management that has been continuous or nearly continuous for a long time (at least 50 years). Site management may have been shorter if management has been prolonged (at least a century or so) before discontinuation and if traditional biotope values are preserved. In both cases, no fertilization or management that reduces traditional biotope values. | No marked overgrowth. | No invasive alien plant species present. |
| **0.9** |  |  |  |  |  |  |
| **0.8** |  |  |  |  |  |  |
| **0.7 (GOOD)** | Vegetation representativeness is decreased to a degree. Essential community characteristic of habitat type is observable but is not as representative as in the ‘excellent’ category. Vegetation structure may be altered to a degree. | Moderate levels of regionally notable traditional biotope plant species and/or a significant site for other notable or endangered biota. |  | Ongoing near-traditional management, or similar, that has continued for a long time, at most, low levels of fertilization or management that decreases site quality. | Intermittent overgrowth. |  |
| **0.6** |  |  |  |  |  |  |
| **0.5 (SATISFACTORY)** | Vegetation representativeness is distinctly decreased. Community and/or vegetation structure exhibit clear changes, some representative species are present. | Some regionally notable traditional biotope plant species and/or other organisms. | Some eutrophication, or moderate levels of problematic species. |  |  | Individual invasive alien plants present. |
| **0.4** |  |  |  |  |  |  |
| **0.3 (POOR)** |  |  |  | Management has discontinued for a long time, or site is under management but management is harmful to traditional biotope values and is not traditional (e.g. fertilization, supplementary feed, overgrazing). | Pronounced overgrowth. | Several occurrences of invasive alien plants. |
| **0.2** |  |  |  |  |  |  |
| **0.1 (VERY POOR)** | Vegetation is unrepresentative. Community is greatly altered, and/or species characteristic of habitat type and site are barely present, vegetation structure is uncharacteristic of habitat type. | No notable traditional biotope plant species or other organisms. | Marked eutrophication, and/or abundant problematic species. | Managament has ended, or site is managed but traditional biotope values have not formed yet, or site has greatly suffered from incorrect management (e.g. fertilization, supplementary feed). | Fully overgrown. | Area is largerly taken over by invasive alien plants. |
| **0.0 (Not a habitat type)** |  | | | | | |

^1^Lists of regionally notable plant species: Kemppainen 2017. Perinnemaisemien inventointiohje. Varsinais-Suomen elinkeino-, liikenne- ja ympäristökeskuksen raportteja 25/2017. In Finnish.

#### 10.3. Wooded traditional biotopes

|  | **(Primary) Vegetation representativeness (vascular plants and/or lichens and/or mosses): community and vegetation structure (abundance, coverage, species/species complex abundance ratios)** | **(Primary)**  **Notable community** | **(Primary)**  **Eutrophication and problematic species** | **(Secondary)**  **Reaping or grazing that does not cause eutrophication, or other comparable management** | **(Secondary)**  **Overgrowth** | **(Secondary)**  **Invasive alien** **plant species** | **(Primary)**  **Representativeness of woodland and scrub^2^**  *Must be distinguished from overgrowth.* |
| --- | --- | --- | --- | --- | --- | --- | --- |
| **Relative attribute weight** | **2** | **2** | **2** | **1** | **1** | **1** | **2** |
| **1.0 (EXCELLENT)** | Representative vegetation. Community and vegetation structure are charcteristic and representative of habitat type, geographical area, and local conditions. | Abundant regionally notable traditional biotope plant species^1^ and/or a significant site for other notable or endangered organisms (e.g. abundant food plants or endangered insects, or knowledge of a viable population of an endangered population). | No marked eutrophication or problematic species. | Ongoing management that has been continuous or nearly continuous for a long time (at least 50 years). Site management may have been shorter if management has been prolonged (at least a century or so) before discontinuation and if traditional biotope values are preserved. In both cases, no fertilization or management that reduces traditional biotope values. | No marked overgrowth. | No invasive alien plant species present. | Forest and scrub are characteristic and representative of habitat type. |
| **0.9** |  |  |  |  |  |  |  |
| **0.8** |  |  |  |  |  |  |  |
| **0.7 (GOOD)** | Vegetation representativeness is decreased to a degree. Essential community characteristic of habitat type is observable but is not as representative as in the ‘excellent’ category. Vegetation structure may be altered to a degree. | Moderate levels of regionally notable traditional biotope plant species and/or a significant site for other notable or endangered biota |  | Ongoing near-traditional management, or similar, that has continued for a long time, at most, low levels of fertilization or management that decreases site quality. | Intermittent overgrowth. |  |  |
| **0.6** |  |  |  |  |  |  |  |
| **0.5 (SATISFACTORY)** | Vegetation representativeness is distinctly decreased. Community and/or vegetation structure exhibit clear changes, some representative species are present. | Some regionally notable traditional biotope plant species and/or other organsims. | Some eutrophication, or moderate levels of problematic species. |  |  | Individual invasive alien plants present. | Forest shows signs of traditional land use, or developing forest and scrub have not become representative yet. |
| **0.4** |  |  |  |  |  |  |  |
| **0.3 (POOR)** |  |  |  | Management has been discontinued for a long time, or site is under management but management is harmful to traditional biotope values and not traditional (e.g. fertilization, supplementary feed, overgrazing). | Pronounced overgrowth. | Several occurrences of invasive alien plants. |  |
| **0.2** |  |  |  |  |  |  |  |
| **0.1 (VERY POOR)** | Unrepresentative vegetation. Community is greatly altered and/or species characteristic of habitat type and site are barely present, vegetation structure is uncharacteristic of habitat type. | No notable traditional biotope plant species or other organisms. | Marked eutrophication, and/or abundant problematic species. | Managament has ended, or site is managed but traditional biotope values have not formed yet, or site has greatly suffered from incorrect management (e.g. fertilization, supplementary feed). | Fully overgrown. | Area is largerly taken over by invasive alien plants. | Forest and scrub are not representative of habitat type, for example due to intensive forestry practices. |
| **0.0 (Not a habitat type)** |  | | | | | | |

^1^Lists of regionally notable plant species: Kemppainen 2017. Perinnemaisemien inventointiohje. Varsinais-Suomen elinkeino-, liikenne- ja ympäristökeskuksen raportteja 25/2017. In Finnish.

^2^Descriptions of trees and scrub representative of wooded traditional biotopes: Kemppainen 2017.

#### 10.4. Alluvial meadows and lake- and rivershore meadows

|  | **(Primary)**  **Vegetation representativeness (vascular plants and/or lichens and/or mosses): community and vegetation structure (abundance, coverage, species/species complex abundance ratios)** | **(Primary)**  **Notable community** | **(Primary)**  **Eutrophication and problematic species** | **(Secondary)**  **Reaping or grazing that does not cause eutrophication, or other comparable management** | **(Secondary)**  **Overgrowth** | **(Secondary)**  **Invasive alien** **plant species** | **(Primary for alluvial meadows, secondary for lake- and rivershore meadows)**  **Flood conditions** | |
| --- | --- | --- | --- | --- | --- | --- | --- | --- |
| **Relative attribute weight** | **2** | **2** | **2** | **1** | **1** | **1** | **Alluvial meadows 2** | **Lake- and rivershore meadows 1** |
| **1.0 (EXCELLENT)** | Representative vegetation. Community and vegetation structure are charcteristic and representative of habitat type, geographical area, and local conditions. | Abundant regionally notable traditional biotope plant species^1^ and/or significant site for other notable or endangered organisms (e.g. abundant food plants of endangered insects, or knowledge of a viable population of an endangered population). | No marked eutrophication or problematic species. | Ongoing management that has been continuous or nearly continuous for a long time (at least 50 years). Site management may have been shorter if management has been prolonged (at least a century or so) before discontinuation and if traditional biotope values are preserved. In both cases, no fertilization or management that reduces traditional biotope values. | No marked overgrowth. | No invasive alien plant species present. | Natural or near-natural flood conditions. | |
| **0.9** |  |  |  |  |  |  |  | |
| **0.8** |  |  |  |  |  |  |  | |
| **0.7 (GOOD)** | Vegetation representativeness is decreased to a degree. Essential community characteristic of habitat type is observable but is not as representative as in the ‘excellent’ category. Vegetation structure may be altered to a degree. | Moderate levels of regionally notable traditional biotope plant species and/or a significant site for other notable or endangered biota. |  | Ongoing near-traditional management, or similar, that has continued for a long time, at most, low levels of fertilization or management that decreases site quality | Intermittent overgrowth. |  |  | |
| **0.6** |  |  |  |  |  |  |  | |
| **0.5 (SATISFACTORY)** | Vegetation representativeness is distinctly decreased. Community and/or vegetation structure exhibit clear changes, some representative species are present. | Some regionally notable traditional biotope plant species and/or other organisms. | Some eutrophication, or moderate levels of problematic species. |  |  | Individual invasive alien plants present. | Flood conditions moderately altered e.g. due to water regulation. | |
| **0.4** |  |  |  |  |  |  |  | |
| **0.3 (POOR)** |  |  |  | Management has discontinued for a long time, or site is under management but management is harmful to traditional biotope values and is not traditional (e.g. fertilization, supplementary feed, overgrazing). | Pronounced overgrowth. | Several occurrences of invasive alien plants.. |  | |
| **0.2** |  |  |  |  |  |  |  | |
| **0.1 (VERY POOR)** | Unrepresentative vegetation. Community is greatly altered, and/or species characteristic of habitat type and site are barely, vegetation structure is uncharacteristic of habitat type. | No notable traditional biotope plant species or other organisms. | Marked eutrophication, and/or abundant problematic species. | Managament has ended, or site is managed but traditional biotope values have not formed yet, or site has greatly suffered from incorrect management (e.g. fertilization, supplementary feed). | Fully overgrown. | Area is largerly taken over by invasive alien plants. | Flood conditions are markedly altered e.g. due to water regulation. | |
| **0.0 (Not a habitat type)** |  | | | | | | | |

^1^Lists of regionally notable plant species: Kemppainen 2017. Perinnemaisemien inventointiohje. Varsinais-Suomen elinkeino-, liikenne- ja ympäristökeskuksen raportteja 25/2017. In Finnish.

### 11. Fell habitats

#### 11.1. Mountain birch forests, Mountain birch scrubs, Mountain forests with aspen

|  | **(Primary)**  **Vegetation representativeness (vascular plants and/or lichens and/or mosses): community and vegetation structure (abundance, coverage, species/species complex abundance ratios)** | **(Primary)**  **Forest structure and regeneration** | **(Secondary)**  **Other anthropogenic impact**  *Does not include the impact of reindeer grazing. However, erosion caused by the reindeer industry in e.g. the feeding and round-up enclosures is included.* |
| --- | --- | --- | --- |
| **Relative attribute weight** | **2** | **2** | **1** |
| **1.0 (EXCELLENT)** | Representative vegetation. Community and vegetation structure are characteristic and representative of habitat type, geographical area, local conditions, and natural successional stage. | A well-regenerating forest. Abundant tree shoots at trunk bases, seedlings/saplings occur regularly, forest has several age classes, and abundant young trees. | No signs or very few signs of anthropogenic activity. At most, individual, narrow paths. |
| **0.9** |  |  |  |
| **0.8** |  |  |  |
| **0.7 (GOOD)** |  | Moderately well regenerating forest. Tree shoots occur regularly, seedlings/saplings are present, young trees can be observed. | Minor anthropogenic activity. For example, abundant paths, ATV tracks. |
| **0.6** |  |  |  |
| **0.5 (SATISFACTORY)** | Vegetation representativeness is decreased to a degree. Essential community characteristic of habitat type is observable but is not as representative as in the ‘excellent’ category. Vegetation structure may be altered to a degree. |  | Fairly abundant anthropogenic activity. For example, terrain erosion, abundant wide paths. |
| **0.4** |  |  |  |
| **0.3 (POOR)** | Vegetation representativeness is distinctly decreased. Community and/or vegetation structure exhibit clear changes, some representative species are present. | Poorly regenerating forest. At most, very few tree shoots, no seedlings/saplings, but young trees are present. |  |
| **0.2** |  |  |  |
| **0.1 (VERY POOR)** | Unrepresentative vegetation. Community is greatly altered, and/or species characteristic of habitat type and site are barely present, vegetation structure is uncharacteristic of habitat type. | Non-regenerating forest. No tree shoots, seedlings/saplings, or young trees are present. Lowest branches have been foraged. | Intensive anthropogenic activity. For example, marked terrain erosion, soil tillage. |
| **0.0 (Not a habitat type)** |  | | |

#### 11.2. Mountain forests with pine, Mountain forests with spruce

|  | **(Primary)**  **Forest structure: uneven-agedness, stratification, stochastic spatial distribution, dead wood present, ancient trees present** | **(Secondary)**  **Vegetation representativeness (vascular plants and/or lichens and/or mosses): community and vegetation structure (abundance, coverage, species/species complex abundance ratios)** | **(Secondary)**  **Other anthropogenic impact**  *Does not include the impact of reindeer grazing. However, erosion caused by the reindeer industry in e.g. the feeding and round-up enclosures is included.* |
| --- | --- | --- | --- |
| **Relative attribute weight** | **2** | **1** | **1** |
| **1.0 (EXCELLENT)** | At least three structural attributes present. | Representative vegetation. Community and vegetation structure are characteristic and representative of habitat type, geographical area, local conditions, and natural successional stage. | No signs or very few signs of anthropogenic activity. At most, individual narrow paths. |
| **0.9** |  |  |  |
| **0.8** |  |  |  |
| **0.7 (GOOD)** |  |  | Minor anthropogenic activity. For example, abundant paths, ATV tracks. |
| **0.6** |  |  |  |
| **0.5 (SATISFACTORY)** | Two structural attributes present. | Vegetation representativeness is decreased to a degree. Community characteristic of habitat type is observable but is not as representative as in the ‘excellent’ category. Vegetation structure may be altered to a degree. | Fairly abundant anthropogenic activity. For example, terrain erosion, abundant wide paths. |
| **0.4** |  |  |  |
| **0.3 (POOR)** | One structural attribute present. | Vegetation representativeness is distinctly decreased. Community and/or vegetation structure exhibit clear changes, some representative species are present. |  |
| **0.2** |  |  |  |
| **0.1 (VERY POOR)** | No structural attributes present. | Unrepresentative vegetation. Community is greatly altered, and/or species characteristic of habitat type and site are barely present, vegetation structure is uncharacteristic of habitat type. | Intensive anthropogenic activity For example, marked terrain erosion, soil tillage. |
| **0.0 (Not a habitat type)** |  | | |

#### 11.3. Other mountain heath scrubs besides mountain birch scrubs; Mountain heaths, Low-graminoid mountain heaths, Mountain meadows, Snowbeds and snow patches

|  | **Vegetation representativeness (vascular plants and/or lichens and/or mosses): community and vegetation structure (abundance, coverage, species/species complex abundance ratios)** | **Other anthropogenic impact**  *Does not include the impact of reindeer grazing. However, erosion caused by the reindeer industry in e.g. the feeding and round-up enclosures is included.* |
| --- | --- | --- |
| **Relative attribute weight** | **1** | **1** |
| **1.0 (EXCELLENT)** | Representative vegetation. Community and vegetation structure are characteristic and representative of habitat type, geographical area, local conditions, and natural successional stage. | No signs or very few signs of anthropogenic activity. At most, individual narrow paths. |
| **0.9** |  |  |
| **0.8** |  |  |
| **0.7 (GOOD)** |  | Minor anthropogenic activity. For example, abundant paths, ATV tracks. |
| **0.6** |  |  |
| **0.5 (SATISFACTORY)** | Vegetation representativeness is decreased to a degree. Essential community characteristic of habitat type is observable but is not as representative as in the ‘exellent’ category. Vegetation structure may be altered to a degree. | Fairly abundant anthropogenic activity. For example, terrain erosion, abundant wide paths. |
| **0.4** |  |  |
| **0.3 (POOR)** | Vegetation representativeness is distinctly decreased. Community and/or vegetation structure exhibit clear changes, some representative species are present. |  |
| **0.2** |  |  |
| **0.1 (VERY POOR)** | Unrepresentative vegetation. Community is greatly altered, and/or species characteristic of habitat type and site are barely present, vegetation structure is uncharacteristic of habitat type. | Intensive anthropogenic activity. For example, marked terrain erosion, soil tillage. |
| **0.0 (Not a habitat type)** |  | |

#### 11.4. Patterned grounds, Solifluction sheets, Frost-influenced heaths

|  | **(Primary)**  **Effects of ground frost** | **(Secondary)**  **Vegetation representativeness (vascular plants and/or lichens and/or mosses): community and vegetation structure (abundance, coverage, species/species complex abundance ratios)** | **(Secondary)**  **Other anthropogenic impact**  *Does not include the impact of reindeer grazing. However, erosion caused by the reindeer industry in e.g. the feeding and round-up enclosures is included.* |
| --- | --- | --- | --- |
| **Relative attribute weight** | **2** | **1** | **1** |
| **1.0 (EXCELLENT)** | Characteristic impact of ground frost in clearly present in habitat type. Wide structural shapes of patterned grounds and solifluction sheets or the unvegetated acrotelm of frost-influenced heaths are present. Herb and/or ground layer closure are barely present. | Representative vegetation. Community and vegetation structure characteristic and representative of habitat type, geographical area, local conditions, and natural successional stage. | No signs or very few signs of anthropogenic activity. At most, individual narrow paths. |
| **0.9** |  |  |  |
| **0.8** |  |  |  |
| **0.7 (GOOD)** |  |  | Minor anthropogenic activity. For example, abundant paths, ATV tracks. |
| **0.6** |  |  |  |
| **0.5 (SATISFACTORY)** | Impact of ground frost is clearly reduced, and herb and/or ground layer closure has begun. Vegetation is fairly widely spread, and the share of unvegetated surface is clearly decreased, but unvegetated surfaces are still present. | Vegetation representativeness is decreased to a degree. Essential community characteristic of habitat type is observable but is not as representative as in the ‘excellent’ category. Vegetation structure may be altered to a degree. | Fairly abundant anthropogenic activity. For example, terrain erosion, abundant wide paths. |
| **0.4** |  |  |  |
| **0.3 (POOR)** |  | Vegetation representativeness is distinctly decreased. Community and/or vegetation structure exhibit clear changes, some representative species are present. |  |
| **0.2** |  |  |  |
| **0.1 (VERY POOR)** | Impact of ground frost has ceased. Herb and/or ground layer is enclosed throughout, woody plant incidence is possibly increased. Only fossil structural shapes are present. | Unrepresentative vegetation. Community is greatly altered, and/or species characteristic of habitat type and site are barely present, vegetation structure is uncharacteristic of habitat type. | Intensive anthropogenic activity. For example, marked terrain erosion, soil tillage. |
| **0.0 (Not a habitat type)** |  | | |
