## Supplement 5 for "Defining ecologically realistic biodiversity offset multipliers with the Response-based Habitat Hectare Assessment of Biodiversity Gains (REHAB)"

**Supplementary 5 – Offset multipliers for restoration actions based on varying initial condition**

We calculated response functions and the corresponding offset action multipliers for restoration actions by iteratively increasing the initial condition from the lowest possible (for the given action) to 0.9 hha/ha by 0.1 interval and capping the yearly condition values to 1 (Figure S5.1.). Multipliers were then calculated separately for each response from the different initial conditions. Thus, the multipliers presented apply to cases where restoration is done in sites higher than the lowest possible condition. We did not modify the response functions otherwise. Multipliers are calculated over the years 0–30 like in the main text.

The distributions of offset multipliers (N=1,551) are shown in Figure S5.2. This distribution follows a similar trend compared to multipliers calculated from the lowest possible initial condition only.

Note that this exercise is done purely to illustrate how applying the initial condition affects offset multipliers for restoration actions, as there are cases where an action cannot be applied in high initial conditions. For example, the action ‘establishing new ponds’ cannot be applied to initial condition other than 0 – otherwise, a *new* pond would not be established. Figure S5.2. includes multipliers for all initial conditions without any ecological considerations regarding the highest appropriate condition for any restoration action. Thus, this figure should be viewed as a demonstration of the general patterns in the offset multiplier distributions rather than a comprehensive set of ecologically applicable multipliers.

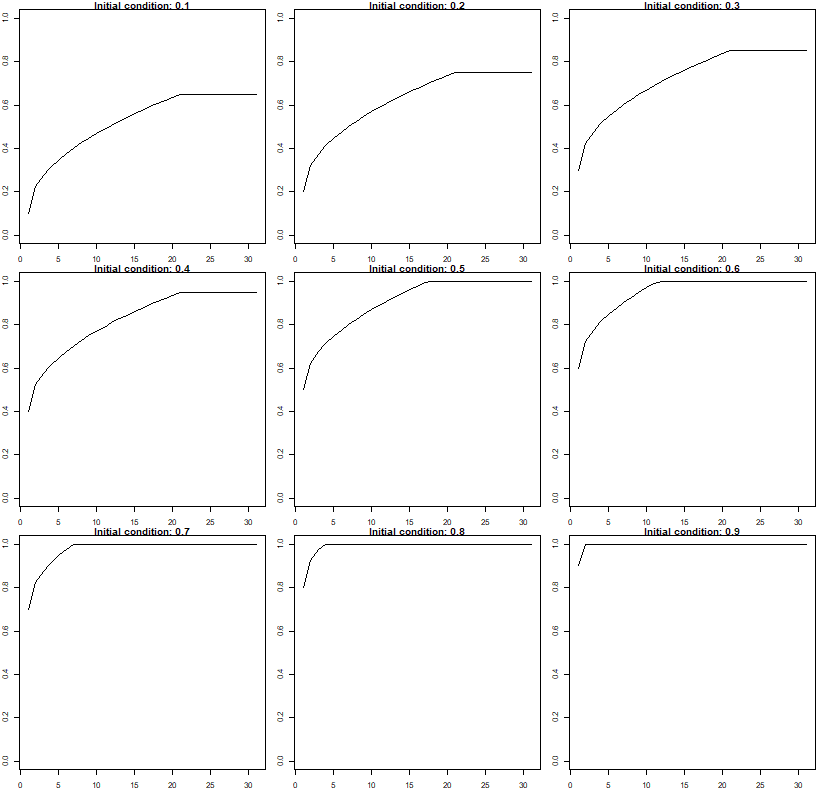

**Figure S5.1.** An example of a restoration response from varying initial conditions. The action improves the target habitat type’s condition from 0.1 to 0.6 hha/ha (y-axis) in 20 years (x-axis). Starting from the initial condition 0.2, the final condition would be 0.7, etc. Yearly response values are capped to 1 to prevent impossibly high condition improvement and thereby too great conservation gains. Thus, when the initial condition in this example exceeds 0.5 hha/ha, the final condition is reached earlier, and the total improvement becomes reduced. From this capped condition improvement function, average gain and the corresponding offset multiplier can be calculated.

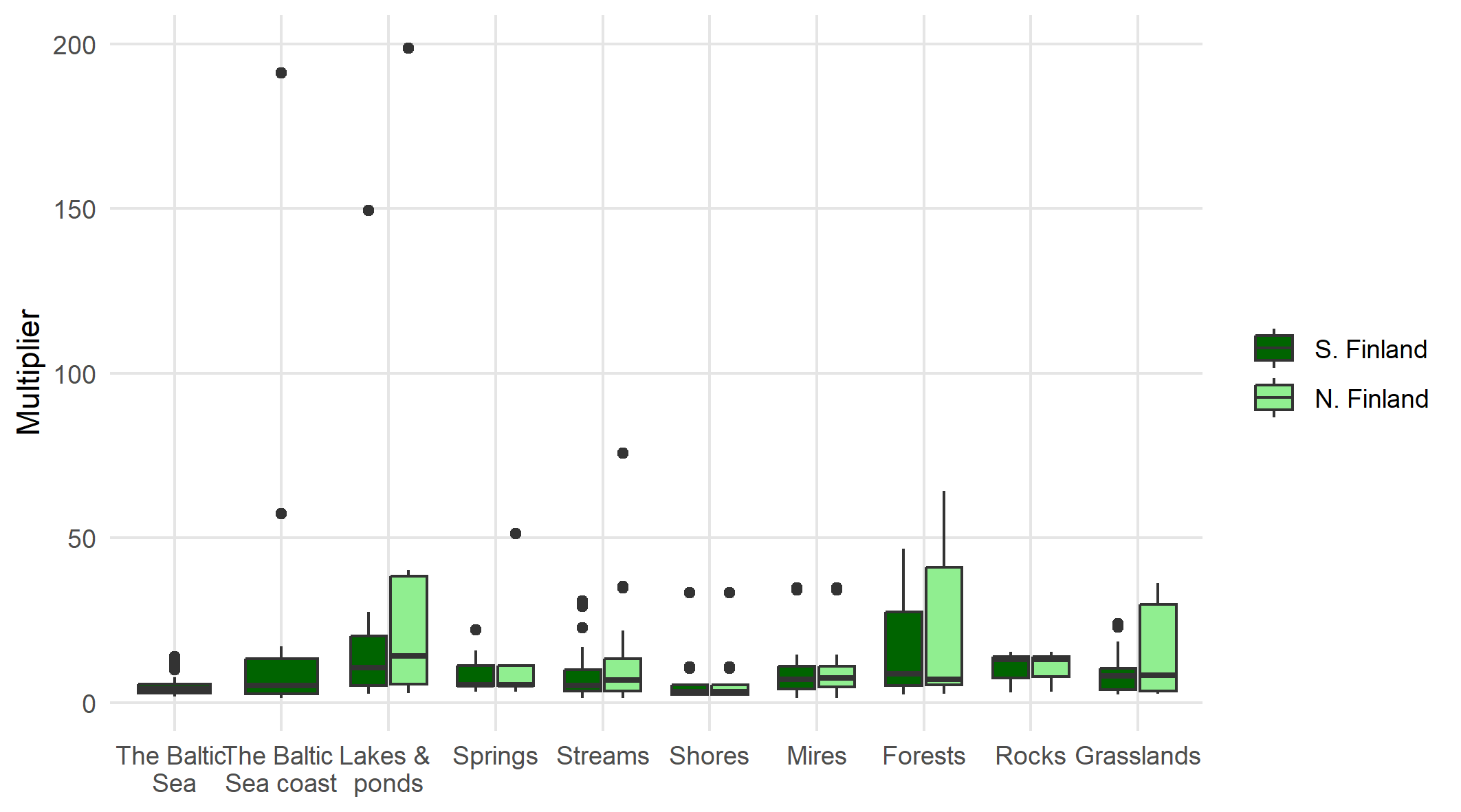

**Figure S5.2.** Distributions of offset multipliers for restoration actions per ecosystem category, when offset multipliers are calculated separately for various initial conditions (from the lowest possible initial condition to 0.9 hha/ha, by 0.1 interval). Refer to Figure 4a in the main manuscript.
